# The emerging fungal pathogen *Candida auris* induces IFNγ to colonize mammalian hair follicles

**DOI:** 10.1101/2025.01.15.632653

**Authors:** Eric Dean Merrill, Victoria Prudent, Parna Moghadam, Abram Rodriguez, Charlotte Hurabielle, Elina K.C. Wells, Pauline Basso, Tiffany C. Scharschmidt, Michael D. Rosenblum, Ari B. Molofsky, Suzanne M. Noble

## Abstract

Public health alarm concerning the emerging fungus *Candida auris* is fueled by its antifungal drug resistance and propensity to cause deadly outbreaks. Persistent skin colonization drives transmission and lethal sepsis although its basis remains mysterious. We compared the skin colonization dynamics of *C. auris* with its relative *C. albicans*, quantifying skin fungal persistence and distribution and immune composition and positioning. *C. auris* displayed a higher propensity to colonize hair follicles and avidly bound to human hair. While *C. albicans* triggered an effective sterilizing type 3/17 antifungal immune response driven by IL-17A/F-producing lymphocytes, *C. auris* triggered a type 1, IFNγ-driven immune response targeting hair follicles. Rather than promoting fungal clearance, IFNγ enhanced *C. auris* skin colonization by acting directly on keratinocytes impairing epithelial barrier integrity and repressing antifungal defense programs. *C. auris* exploits focal skin immune responses to create a niche for persistence in hair follicles.

## Main Text

First described in 2009, the fungal pathogen *Candida auris* is now endemic among humans across the globe^1–3^. The emergence of *C. auris* has raised public health alarm because of its high frequency of antifungal drug resistance, which includes pan-resistant isolates^4,5^, and its ability to cause infectious outbreaks in hospitals and long-term care facilities^6,7^. Epidemiologic investigations of outbreaks have identified asymptomatic skin colonization as a key factor fostering transmission and progression to invasive disease.^8–10^ Skin colonization typically persists for many months, and colonized individuals are at high risk for subsequent bloodstream infections,^11,12^ which are associated with high mortality, ranging between 30%-70%^7^. There are currently no effective methods to eradicate *C. auris* from the skin^9^.

Mammals have co-evolved with fungi, including yeasts such as *Malassezia* and *Candida albicans* that colonize human skin^13^. To persist on skin, fungi must establish residence and proliferate while avoiding sterilizing host immunity. Based on studies of *C. albicans,* type 3/17 immune responses, characterized by the cytokines IL-17A, IL-17F, and IL-22, are thought to be the critical effectors of anti-fungal immunity^14^. In humans and mice, mutations and autoantibodies affecting the IL-17A/F signaling pathway render hosts susceptible to *C. albicans* overgrowth and pathology, particularly at barrier sites such as the mouth, genitourinary tract, and nails, a syndrome known in humans as chronic mucocutaneous candidiasis (CMC)^15–17^.

Acting on multiple cell types, type 3/17 cytokines recruit neutrophils and drive epithelial cell proliferation and production of anti-microbial peptides (AMP), ultimately leading to the clearance of extracellular fungi and bacteria^18^. If barrier containment fails and *C. albicans* enters the bloodstream, IFNγ-driven type 1 immunity serves as a critical backup to inhibit fungal sepsis^19–22^. However, excessive IFNγ signaling in the oral epithelium can lead to overgrowth of *C. albicans*, as observed in patients with genetic defects in the AIRE immune tolerance gene^23,24^, suggesting that lymphocyte-driven type 1 and type 3/17 immune responses may function antagonistically, at least in some barrier sites and clinical contexts.

To identify the mechanisms governing fungal interactions with skin, we performed systematic comparisons of *C. auris* and *C. albicans* in mouse models of skin colonization, monitoring both fungal behavior and host immune responses. *C. auris* consistently outperformed *C. albicans* in tests of fungal abundance and persistence and exhibited unique hair-binding activity and a strong tropism to hair follicles. Moreover, whereas *C. albicans* provoked type 3/17, IL-17-driven responses that resulted in fungal clearance, *C. auris* provoked a type 1-dominant, IFNγ response that supported its persistence in hair follicles.

### *Candida auris* triggers a type 1 immune response in the skin

#### *C. auris* colonizes skin with higher titers than *C. albicans* and preferentially resides in hair follicles

To compare the kinetics of *C. auris* and *C. albicans* skin association, we adapted an established model of fungal skin colonization^25^. Yeast was applied to the shaved but undamaged back skin of C57BL/6 mice, followed by an analysis of fungal titers in skin recovered at predefined time points (**Fig1A**). Two days following a single topical application of *C. auris* or *C. albicans*, *C. auris* titers slightly exceeded those of *C. albicans*; however, by day 5, *C. albicans* was no longer detectable in skin homogenates, whereas *C. auris* persisted throughout the 30-day time course (**Fig1B**). Following four serial doses over six days, *C. albicans* was still undetectable by 2 days after the final inoculum whereas *C. auris* persisted in all animals throughout the time course (**Fig1C**). Next, we performed histological analysis to determine fungal localization in colonized skin. After two days of colonization, clusters of *C. auris* yeasts were located at the skin surface and associated with hair follicles, where they were distributed adhering to hair shafts, the follicle orifice, and the hair follicle lumen (**Fig1D-E; Fig1s1A-B**). Conversely, *C. albicans* yeast were associated with the skin surface, with sparse examples of hair follicle localization **(Fig1s1B)**. By day 14, *C. albicans* was no longer identified in the skin, but clusters of *C. auris* could still be identified in hair follicles, primarily localized to the follicle orifice (**Fig1s1C**). We conclude that *C. auris* colonizes mouse skin with greater abundance and greater persistence than *C. albicans,* consistent with other observations^26^,Prudent et al. submitted.

**Figure 1:**
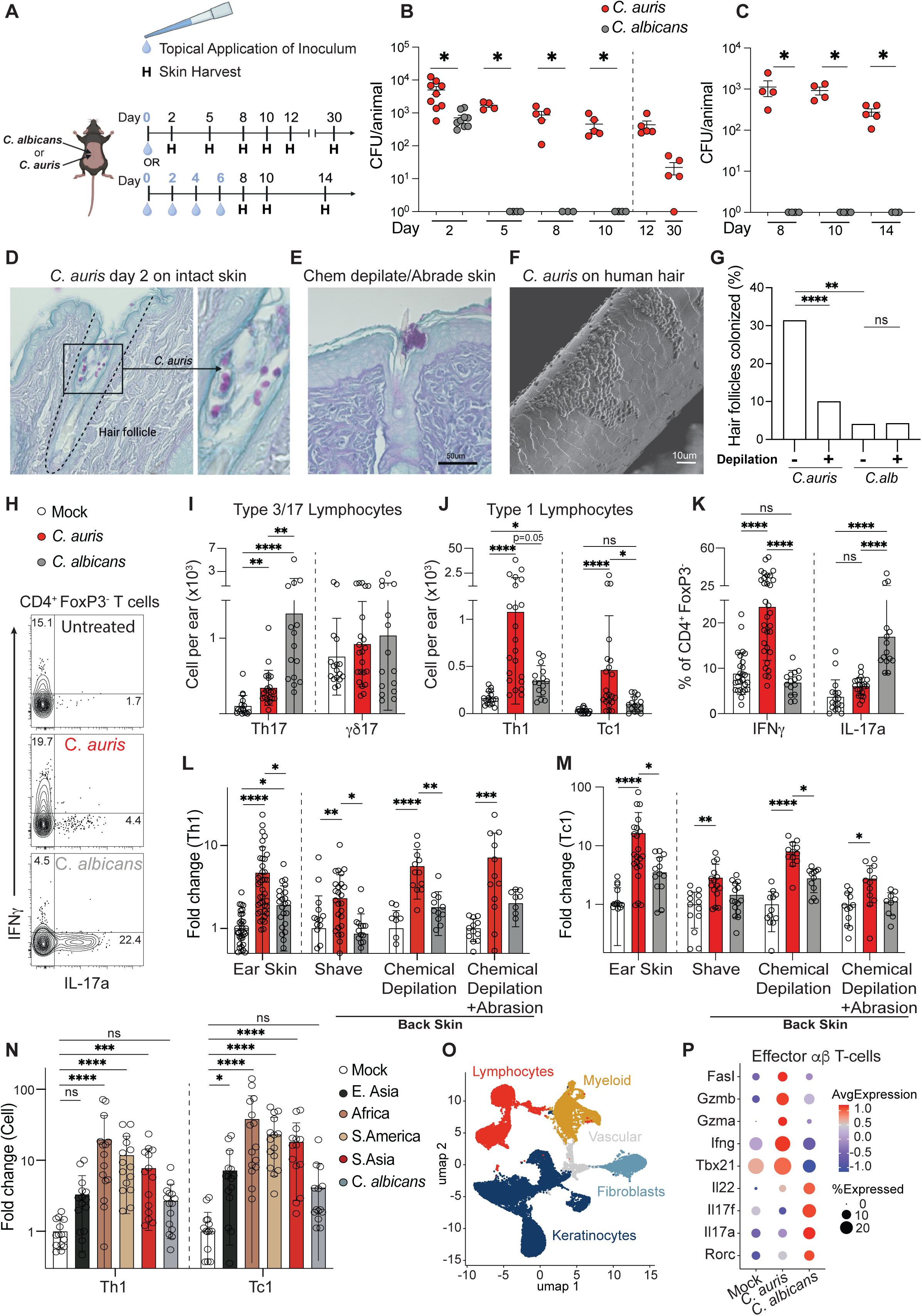
*C. auris* skin and hair follicle colonization elicits a type 1 immune response. **(A)** Schematic of the models for skin topical colonization. For fungal load, C57BL/6 wild-type mouse back skin was harvested using either a single dose (top) or repetitive dosing (bottom) regimen. For flow immunophenotyping, both back and ear skin were harvested. **(B)** *C. auris* or *C. albicans* were topically applied at 10^8^ colony-forming units (CFU) on day 0, and back skin was harvested for CFU on the days indicated. **(C)** *C. auris* or *C. albicans* was topically applied at 10^8^ CFU on days 0,2,4,6 and back skin was harvested on days indicated for CFU. **(D)** *C. auris* colonized skin was stained with Periodic Acid-Schiff (PAS) on day 2, with yeast (dark purple) identified within the hair follicle. **(E)** *C. auris* was visualized in the hair follicle via PAS stain 2 days after epicutaneous infection. **(F)** Scanning electron microscopy image of *C. auris* binding to human hair shaft in vitro. **(G)** *C. auris* or *C. albicans* was colonized on back skin and harvested on day 2, fixed, embedded in paraffin, and stained for PAS. Quantification of hair follicles affected (defined by >1 yeast with hair follicle) for *C. auris* and *C. albicans* +/- chemical depilation with depilating agent Nair^TM^. **(H)** Flow cytometry plot of effector T cells (CD4^+^FoxP3^-^) 14 days after topical colonization with *C. auris* or *C. albicans* (days 0,2,4, and 6). **(I)** Flow quantification of skin Th17 (CD3^+^CD4^+^IL-17a^+^) or γδ17(CD3^+^TCRγδ^+^IL-17a^+^) T cells on D14. **(J)** Flow quantification of skin Th1 (CD3^+^CD4^+^IFNγ^+^) or Tc1 (CD3^+^CD8^+^IFNγ^+^) T cells on D14. **(K)** Percent cytokine-producing (IL-17a and IFNγ) cells of effector T-cells (CD3^+^CD4^+^FoxP3^-^). **(L,M)** Fold change normalized to same experiment controls of (**L**) Th1 and (**M**) Tc1 in mice treated with various levels of epithelial damage. Ear and back skin (Shave) were colonized (days 0,2,4,6) in the absence of inflammation, back skin of mice was depilated and either treated with sandpaper (Depilation+SP) or not (Depilation) prior to application of *C. albicans* or *C. auris* on days 0,2,4,6. **(N)** Th1 and Tc1 numbers as fold change normalized to controls from the same experiment, comparing strains from different clades of *C. auris*. **(O)** UMAP clustering of major cell types from single-cell RNA sequencing of mouse back skin at day 14 following topical colonization with either *C. auris*, *C. albicans*, or mock-colonized controls. **(P)** Expression dotplot of IL-17- and cytotoxicity-associated scRNAseq genes after subsetting on T-cells. *P<0.05, **P<0.01, ***P<0.001, ****P<0.0001 in student T-Test **(B,C)**, Kruskal-Wallis test with Dunn’s multiple comparison test **(I,J,L,M,N)**, one-way analysis of variance (ANOVA) and Tukey’s multiple comparison test **(K)**. PAS = Periodic acid–Schiff. All images are representative of 3 or more experiments.

The histological analysis suggested that *C. auris* can bind directly to hair. To directly test this hypothesis, we developed an *in vitro* hair-binding assay. *C. auris* and *C. albicans* were separately incubated with human hair fragments, stained with calcofluor white, and quantified for adherence using fluorescence microscopy. As predicted, *C. auris* exhibited higher *in vitro* hair-binding activity than *C. albicans* (**Fig1F**; **Fig1s1D,E**). To determine whether the presence of hair facilitates *C. auris* localization to hair follicles, we quantified the fraction of yeast-associated hair follicles in animals colonized using our standard skin colonization protocol or following pre-treatment of skin with chemical depilatory cream (Nair^TM^) to remove the hair stubble that remains after shaving. Pretreatment with depilatory cream reduced the fraction of hair follicles associated with *C. auris* from ∼30% to ∼10% but had no effect on the <5% of follicles associated with *C. albicans* (**Fig1G**). These results indicate that direct binding to mammalian hair promotes the localization of *C. auris* to hair follicles.

#### *C. auris* and *C. albicans* induce distinct immune responses in colonized skin

Type 3/17 immune responses inhibit the proliferation of *C. albicans* at epithelial barrier sites, whereas type 1 responses limit systemic infections. Far less is known about immune defenses against *C. auris,* particularly in skin. We therefore sought to characterize lymphocyte composition and cytokine expression in mouse skin after colonization with *C. auris,* using *C. albicans* for comparison. Four inocula of either yeast species or PBS alone (“mock”) were applied to the ear or shaved back skin of C57BL/6J mice over six days, and the skin was recovered for flow cytometry on day 14. Relative to mock-colonized animals, *C. albicans* stimulated a >10-fold increase in the abundance of Th17 cells and a >3-fold increase in intracellular IL-17A production, consistent with other observations^27^. By comparison, *C. auris* induced relatively blunted Th17 responses and neither fungal species induced significant changes in the abundance or IL-17A production of innate-like γδ T cells at this day 14 time point **(Fig1H-I**). Surprisingly, *C. auris* but not *C. albicans* induced robust increases in the abundance of type 1-polarized lymphocytes, including cytotoxic CD8^+^ Tc1 cells, helper CD4^+^ Th1, and their production of IFNγ (**Fig1J-K**). Finally, neither fungal species affected the abundance of innate or adaptive type 2 lymphocytes (ILC2, Th2), and both species drove minor and variable elevations in regulatory T lymphocytes (Treg; **Fig1s1F,G**).

We next asked whether skin immune responses vary as a function of body site, comparing ear and back skin, or degree of preexisting skin injury, comparing undamaged skin with skin damaged by chemical depilation +/- sandpaper abrasion. *C. auris* elicited increases in Th1 and Tc1 cell abundance and intracellular IFNγ production in all animals, regardless of body site or skin damage (**Fig1L,M**). However, whereas *C. auris* provoked small increases in Th17 abundance in the undamaged ear and back skin, it induced larger increases in damaged skin that were comparable to those induced by *C. albicans* (**Fig1s1H)**. These findings suggest that *C. auris* induces type 1 immune responses to a similar extent in the skin at different body sites, independent of underlying skin injury, whereas the size of the provoked type 3/17 response increases as a function of skin damage.

The preceding experiments used a *C. auris* clinical isolate from Clade I (South Asia). To determine whether diverse *C. auris* strains share the ability to induce type 1 immune responses in skin, we tested clinical isolates representing three other clades, as well as the original *C. auris* strain and *C. albicans.* All four *C. auris* strains provoked increases in type 1 lymphocytes, with the Clade III (Africa) and Clade IV (South America) strains stimulating the strongest responses and the Clade II strain (East Asia) the weakest (**Fig1N**). The different *C. auris* strains also elicited increases of variable size in Th17 abundance, while having a less consistent effect on γδ T cells that produce IL-17A (γδ17 lymphocytes; **Fig1s1I**). We conclude that *C. auris* and *C. albicans* are both capable of inducing type 3/17 lymphocyte responses in skin, whereas *C. auris* drives type 1 immunity.

To capture host responses using an orthogonal approach, we turned to single-cell RNA sequencing (scRNAseq). Wild-type C57BL/6J (WT) mice were colonized with *C. auris*, *C. albicans*, or PBS alone (mock-colonized) before harvesting skin on day 14 for scRNAseq. Based on the expression of cell type-specific markers, gene expression clusters were mapped to immune lymphocytes and myeloid cells, epithelial keratinocytes, vascular endothelial cells, and stromal fibroblasts based on the expression of cell type-specific markers (**Fig1O; Fig1s2A,B**). We initially focused on transcriptional responses in adaptive T lymphocytes (see **Fig1s2C** for clustering), where upregulation of type 3/17-associated genes, including Il17a, Il17f, Il22, and the transcription factor Rorc, was apparent in response to *C. albicans* skin colonization, but not *C. auris* (**Fig1P**). Rather, exposure to *C. auris* was associated with the upregulation of type 1-associated immune genes, including the transcription factor Tbet (Tbx21), Ifng, and cytotoxicity-associated factors (Fasl, Gzma, Gzmb), in adaptive T cells (**Fig1P**). Consistent with our flow cytometric results, the scRNAseq data suggest that *C. auris* skin colonization drives a type 1, IFNγ-associated immune response that is distinct from the response to *C. albicans*.

### *C. auris* induces a type 1 immune niche in the upper hair follicle

#### Type 1 and type 3/17 lymphocytes occupy distinct epidermal and dermal niches

Reflecting its function as a barrier tissue, skin is partitioned between the microbial-dense and epithelial cell-rich epidermis and the underlying dermis. The epidermis contains hair follicles composed of epithelial cells (keratinocytes) and associated sebaceous glands, while the dermis houses sensory and autonomic neurons, blood vessels, lymphatics, and fibroblast and adipocyte support cells (**Fig2A**). Hair follicles are important for skin immunology, functioning both as reservoirs of epithelial stem cells that mediate skin homeostasis and repair and as direct portals to the microbe-dense external environment^28^. To create a portrait of skin lymphocyte topography, we used volumetric confocal microscopy with tissue clearing.

**Figure 2:**
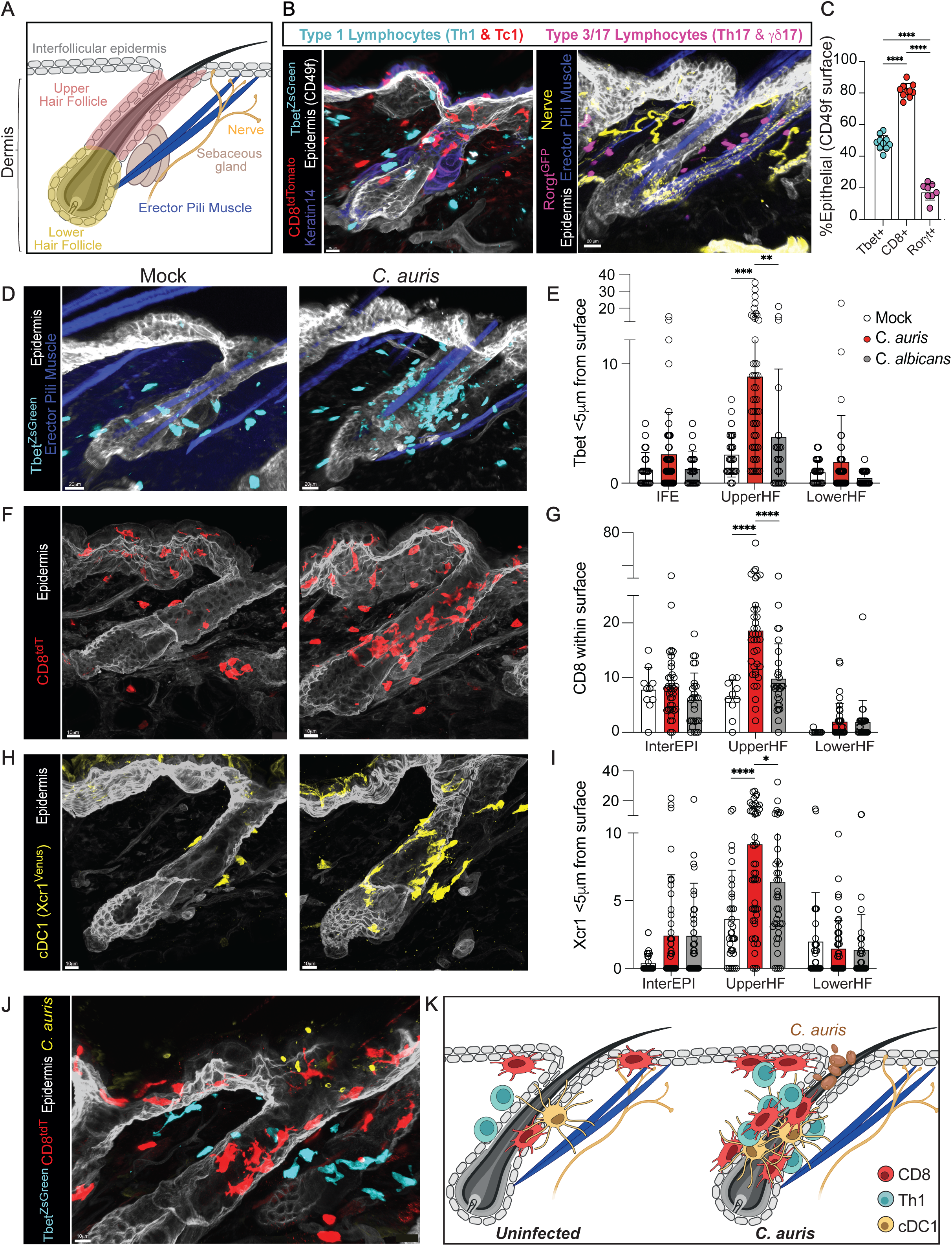
*C. auris* expands a type 1 immune niche in the upper hair follicle. **(A)** Schematic of the skin ‘topography’, including the dermis and the epidermis, itself composed of keratinocytes that also line hair follicles. **(B)** T cell subsets in the back skin were visualized using thick section confocal microscopy with tissue clearing of untreated adult mice. Left panel: type 1 lymphocytes (Th1 defined as Tbet^ZsGreen^ and Tc1 defined as CD8^Cre^;R26^TdT+^). Right panel: type 3/17 lymphocytes defined as Rorgt^GFP^. **(C)** Quantification of intra-epithelial lymphocytes. An epithelial cell surface was defined using CD49f and immune cells were quantified as intraepithelial if their distance was less than 0um to the surface. Each dot is an average of numerous spots from a large image collected at 20x. **(D-I)** Tbet^ZsGreen^, CD8^+^ cytotoxic lymphocytes (CD8^Cre^;R26^TdT+^), or cDC1 (Xcr1^Venus^) were quantified in relation to the epidermis (CD49f/Itga6 stain). Using the ‘Spot’ feature in Imaris, CD8 spots were defined as within an epithelial region when <0um from the respective surface and Tbet and cDC1 spots were defined as near a surface when <5um from the respective surface. Each dot represents a single hair follicle taken from 3-15 hair follicles per mouse and from at least 3 mice per group. **(J)** Thick-section confocal image using anti-Candida antibody, showing clusters of *C. auris* in proximity to intraepithelial CD8^+^ T-cells with type 1 Tbet^ZsGreen^ lymphocytes predominantly in the nearby dermis. **(K)** Graphical representation of the skin immune cell positioning, with *C. auris* encompassed by intraepithelial CD8^+^ T cells, and peri-follicular cDC1 and Th1 CD4^+^ T cells. Images are representative of 3 or more experiments and were from mice with hair follicles in their resting telogen cycle (∼8-10 weeks old). All mice labeled *C. auris* or *C. albicans* were treated on days 0,2,4, and 6 and back skin was harvested on day 14 **(E-J)**. *P<0.05, **P<0.01, ***P<0.001, ****P<0.0001 for one-way analysis of variance (ANOVA) and Tukey’s multiple comparison test **(C,G,I).**

We created a map of skin immune cell topology, starting with specific pathogen-free (SPF) wild-type mice that were not colonized with fungi. Type 1 and type 3/17 lymphocytes were visualized using mice that express fluorescently labeled reporters for the lymphocyte program-specific transcription factors, Tbet^ZsGreen^ and Rorγt^GFP^, respectively; epithelial cells, neurons, and erector pili muscles were stained with specific fluorescent antibodies. Type 1 lymphocytes (Tbet^ZsGreen^), including both Tc1 and Th1 cells, were observed to be relatively evenly distributed between the epidermis and dermis. Using mice that express a reporter for CD8^+^ T cells (CD8^Cre^; R26^tdT^), Tc1 lymphocytes (CD8^+^ Tbet^+/-^) were distinguished from Th1 cells and found to be predominantly (∼80%) localized within the epithelium (**Fig2B,C, Fig2s1A,B**). In contrast, type 3/17 lymphocytes (Rorγt^GFP^) were localized primarily in interfollicular (between hair follicles) regions of the upper dermis, often near erector pili muscles and their associated neurons (**Fig2B,C, Fig2s1C,D)**. To more precisely map type 1 lymphocytes within the epithelium, we counted the number of cells localized to the interfollicular epidermis (at the skin surface), the upper hair follicle, or the lower hair follicle **(Fig2A, Fig2s1E)**. Overall, type 1 lymphocytes (Tbet^+^) were primarily localized near the upper portion of hair follicles **(Fig2D,E)**, with CD8^+^ T cells distributed between the upper hair follicle and the interfollicular epidermis (**Fig2F,G)**.

#### Antigen-presenting cells are localized near their cognate lymphocytes

Antigen-presenting cells (APC) orchestrate adaptive immune responses in lymph nodes and tissues. In mouse skin, the main types of APC are Langerhans cells (LC), type 1 and type 2 conventional dendritic cells (cDC1 and cDC2), and monocyte-derived dendritic cells (MoDC)^29^. We hypothesized that the topography of dendritic cell subtypes might mirror the resting distribution of related type 3/17 and type 1 lymphocytes. Indeed, cDC2 (Mgl2^Cre^; R26^tdTomato^, which also labels some macrophages; dermal CD11c^Cre^; R26^tdTomato^) were visualized throughout the dermis **(Fig2s2A-D)**, similar to type 3/17 lymphocytes (**Fig2B-C)**, whereas cDC1 (Xcr1^Venus/+^) were positioned near the epithelial surface (<5um). Unlike epidermal-resident Langerhans cells (intraepithelial, ramified CD11c^Cre^; R26^tdTomato^) (**Fig2s2B**)^30^, cDC1 were not embedded within the epithelium proper but, rather, enriched around the upper hair follicle and adjacent to the epithelial basement membrane (**Fig2s2E-F**). We conclude that skin cDC1 and the type 1 lymphocytes are closely approximated in an upper hair follicle niche, with sub-epithelial Th1 cells poised to reinforce the actions of intraepithelial CD8^+^ Tc1. In contrast, cDC2 and type 3/17 lymphocytes are preferentially enriched in the dermis.

#### *C. auris* promotes the expansion of type 1 immune cells near the upper hair follicle

Next, we assessed the impact of fungal skin colonization on the distribution of lymphocytes and APC. Following association with *C. auris* but not *C. albicans*, increased type 1 lymphocytes were observed within and abutting the epithelium of the upper hair follicle **(Fig2E-G, Fig2s1F**). Maintaining similar localization as in mock-colonized skin, type 1-polarized lymphocytes (Tbet^+^) in *C. auris-*colonized skin were located within and slightly deep to the epithelium, with CD8^+^ T cells primarily within the epithelium **(Fig2E,G)**. By day 14, increased cDC1 accumulated near the upper hair follicle of *C. auris-*colonized skin (**Fig2H,I, Fig2s1G**), consistent with the increased abundance of cDC1 detected by flow (**Fig2s2G,H)**. In skin colonized with fluorescently labeled *C. auris,* we observed examples of yeast at the hair follicle orifice, encircled by type 1 lymphocytes and cDC1, as well as examples of cDC1 extending cell projections between hair follicle epithelial cells and in proximity to type 1 lymphocytes **(Fig2J,K**, **Fig2s2I,J)**.

We reviewed the scRNAseq datasets for myeloid cells to determine the impact of *C. auris* and *C. albicans* on skin APC. Myeloid cell clusters mapped to multiple subsets of APC, including LC, cDC1, and cDC2, as well as macrophage subsets **(Fig2s3A)**. In skin colonized with *C. auris,* we observed increased expression of cDC1 markers (e.g. Xcr1, Batf3), whereas *C. albicans* triggered the upregulation of LC markers *(e.g.,* EpCAM, Cdh1). *C. auris* also induced pathways associated with ‘antigen presentation through MHC’ and ‘Stat1 signaling’, as well as MHCI-associated genes (B2m, H2-D1, H2-K1), intracellular antigen processing genes (Tap1, Tap2), and IFNγ stimulated genes (Stat1, Irf1/8/9, Cxcl9; **Fig2s3B**). To determine whether cDC1 contribute to the *C. auris*-associated type 1 immune response, we depleted cDC1 throughout the period of *C. auris* colonization using Xcr1^DTR^ mice. *C. auris* colonization of cDC1-deficient animals produced a blunted type 1 response, with reduced abundance and intracellular IFNγ production by skin Th1 and Tc1 relative to cDC1-replete animals (**Fig2s3C-F**). We conclude that cDC1 cells, which are localized at the upper hair follicle near CD8^+^ cytotoxic T cells and CD4^+^ Th1 cells, are required for the expansion of Tc1 and Th1 cells in response to *C. auris*.

### *Candida albicans* induces a sterilizing IL-17-mediated immune program in the upper hair follicle epithelium

Having established that *C. albicans* and *C. auris* elicit distinct skin immune responses, we investigated the consequences of these lymphocyte-driven immune responses on fungal colonization, beginning with *C. albicans*. By day 14 of the repetitive colonization model, *C. albicans* was completely cleared from the skin of wild-type mice; however, IL-17A/F-deficient animals maintained a high (>10^5^ CFU/animal) fungal burden and substantial skin pathology (**Fig3A,B**), confirming the critical role of IL-17 in anti-Candida barrier immunity^16,19,31,32^. In addition to forming a physical barrier, keratinocytes serve as important immune sensors and effectors of IL-17 signaling^33,34^. To determine whether the anti-Candida function of IL-17 is mediated through keratinocytes, we generated mice that lack the IL-17 receptor specifically in keratinocytes (Ker14^Cre^; IL17Ra^F/F^). Repetitive colonization of Ker14^ΔIL17Ra^ mice phenocopied the colonized IL-17^-/-^ animals (**Fig3A-B**), indicating that IL-17 signaling through keratinocytes is required to restrict the growth of *C. albicans* on the skin.

**Figure 3:**
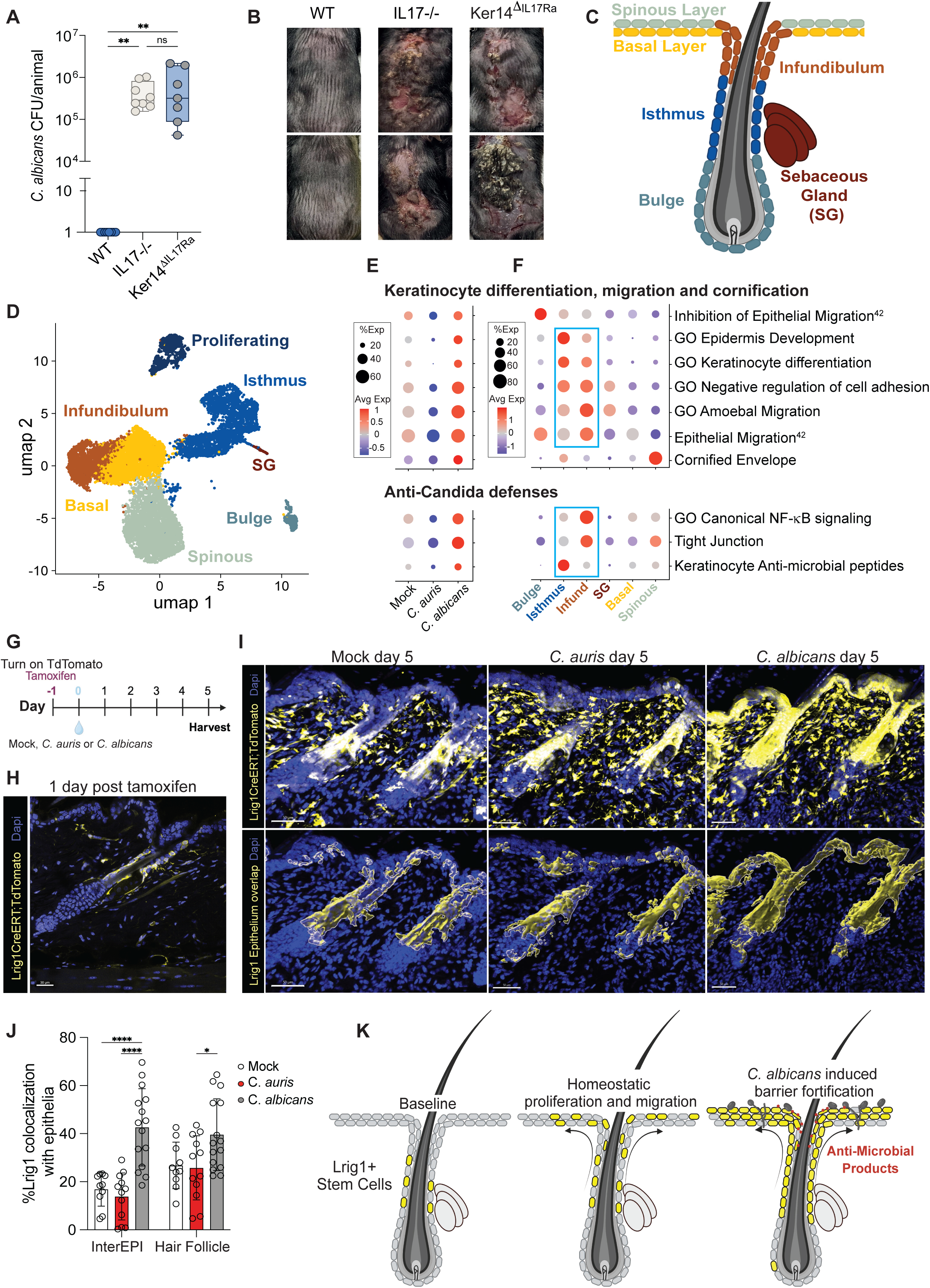
IL-17-mediated sterilizing immunity to *Candida albicans* is directed at the upper hair follicle epithelium. **(A)** Mice were colonized on days 0,2,4,6 with *C. albicans* and back skin was harvested for CFU on day 14 (8 days after the final application). Mice that lack IL-17 receptor on keratinocytes (Ker14^Cre^;IL17Ra^F/F^, i.e. Ker14^ΔIL17Ra^) were colonized alongside littermate controls (IL17Ra^F/F^ lacking Cre recombinase) and IL-17-/- mice, which globally lack IL-17a and IL-17f. **(B)** Representative photographs of back skin from Ker14^ΔIL17Ra^ and littermate controls at day 14. **(C)** Schematic cartoon showing keratinocyte subsets. **(D)** From scRNAseq, keratinocytes were pre-selected and further analyzed to reveal subset clusters: hair follicle bulge, isthmus, infundibulum, basal layer, spinous layer, sebaceous gland (SG), and proliferating. **(E,F)** scRNAseq dot plot by **(E)** treatment condition or **(F)** cell type, showing modules upregulated following *C. albicans* colonization associated with epithelial stem cells, differentiation, migration and cornification associated modules and anti-Candida defenses. **(G)** Schematic showing experimental design where Lrig1^CreERT^; R26^Tdt^ mice were injected with tamoxifen 1 day prior to mock, *C. auris* or *C. albicans* colonization. Mice were harvested for confocal microscopy on day 5 post colonization. **(H)** Representative image from slices of 3D image showing lineage traced Lrig1+ cells 1-day post tamoxifen **(I)** Top panel showing representative 3D image of lineage traced Lrig1+ cells 5 days post-colonization with mock, *C. auris,* or *C. albicans*. Bottom panel showing overlapping Lrig1 and CD49f epithelial surface, using Imaris imaging software. **(J)** Graphical representation of overlapping Lrig1^+^ surface with the CD49f^+^ epithelia surface subdivided into the hair follicle and interfollicular epidermis, using Imaris imaging software. **(K)** Schematic showing proposed *C. albicans* induced changes to keratinocytes. *P<0.05, **P<0.01, ****P<0.0001 for one-way analysis of variance (ANOVA) and Tukey’s multiple comparison test **(A,I).**

To further explore the role of keratinocytes in antifungal defense, we analyzed the scRNAseq data for keratinocytes in mock-colonized, *C. albicans-*colonized, and *C. auris-*colonized skin (**Fig3C-D**, **Fig3s1A**). Pathway analysis of keratinocytes revealed that genes associated with epithelial cell maturation and barrier restoration were upregulated in response to *C. albicans* but downregulated in response to *C. auris*, with specific elements of programs regulated in discrete keratinocyte subtypes (**Fig3E**, **Fig3s1B,C)**. For example, keratinocytes of the hair follicle isthmus exhibited upregulation of epithelial differentiation programs. Keratinocytes of the upper hair follicle (isthmus and infundibulum) exhibited upregulation of genes associated with the inhibition of cell adhesion, a key step for cell migration, and additional cell migration modules were upregulated in the infundibulum. Upper hair follicle keratinocytes also exhibited upregulation of modules associated with anti-*Candida* immunity, such as canonical NFκB signaling, tight junction formation, and anti-microbial product expression (**Fig3F**).

We were intrigued by the finding that *C. albicans* but not *C. auris* induces keratinocytes of the upper hair follicle—a region that functions as an organizational hub for skin immune cells^28^—to upregulate genetic programs that support epithelial barrier function. To validate these findings, we utilized mice that label upper hair follicle stem cells with TdTomato following tamoxifen induction (Lrig1^CreERT^; R26^tdT^), enabling the precise tracking of Lrig1^+^ stem cells and their progeny over time (**Fig3G**). One day after tamoxifen treatment, Lrig1^CreERT^ R26^tdT^ animals were inoculated with *C. auris*, *C. albicans,* or PBS (mock), and skin was harvested for imaging after five days of colonization. Skin from *C. albicans-*colonized animals exhibited a greater number of Lrig1^+^ stem cells (TdT-labeled) in the upper hair follicle, as well as more examples of Lrig1^+^ cell migration to the interfollicular dermis than skin from either mock-treated or *C. auris-*colonized animals (**Fig3H-J**). These results suggest that exposure to *C. albicans* increases the proliferation and differentiation of Lrig1^+^ stem cells to enhance epithelial barrier function, whereas *C. auris* does not detectably alter the homeostatic replenishment of epithelial cells in skin (**Fig3H-K**).

### *C. auris* induces IFNγ-mediated epithelial rewiring that promotes skin colonization

Given that (1) the IL-17 anti-*Candida* response is directed at upper hair follicle keratinocytes, (2) *C. auris* colonizes hair follicles, and (3) *C. auris-*induced type 1 lymphocytes surround the upper hair follicle, we hypothesized that keratinocytes of the upper hair follicle may also be targets of IFNγ signaling and play a role in *C. auris* colonization and persistence. Therefore, we mined our scRNAseq datasets for evidence of IFNγ signaling in this cell type. In addition to the previously described transcriptional responses **(Fig3E,F)**, upper hair follicle keratinocytes from *C. auris*-colonized skin exhibited upregulation of multiple canonical IFNγ signaling targets, including Stat1 and MHC class 1 antigen presentation genes, as well as pathways associated with endoplasmic reticulum stress, autophagy, cellular respiration, and necroptosis **(Fig4A,B, Fig4s1A,B)**. To determine whether IFNγ levels affect the transcriptional responses of keratinocytes to *C. auris,* we colonized animals that either lack the IFNγ gene (IFNγ^-/-^) or constitutively overexpressed IFNγ (Yeti/+ mice^35,36^, henceforth referred to as IFNγ^OE^), as well as IL-17^-/-^ and WT mice. After 14 days, skin was recovered for scRNAseq. Consistent with a role for IFNγ signaling on keratinocyte gene expression, we observed an even more dramatic upregulation of IFNγ response and stress genes in the upper hair follicle keratinocytes of *C. auris-*colonized IFNγ^OE^ animals than in WT animals, whereas the same genes were downregulated in IFNγ^-/-^ animals (**Fig4C**). Conversely, barrier integrity and anti-Candida defense pathways were strongly downregulated in keratinocytes from colonized IFNγ^OE^ mice but upregulated in colonized IFNγ^-/-^ mice (**Fig4C, Fig4s1C-E**). Keratinocytes from *C. auris*-colonized IL-17-deficient mice exhibited intermediate gene expression phenotypes. These results support a role for IFNγ in driving *C. auris-*associated responses in hair follicle keratinocytes.

**Figure 4:**
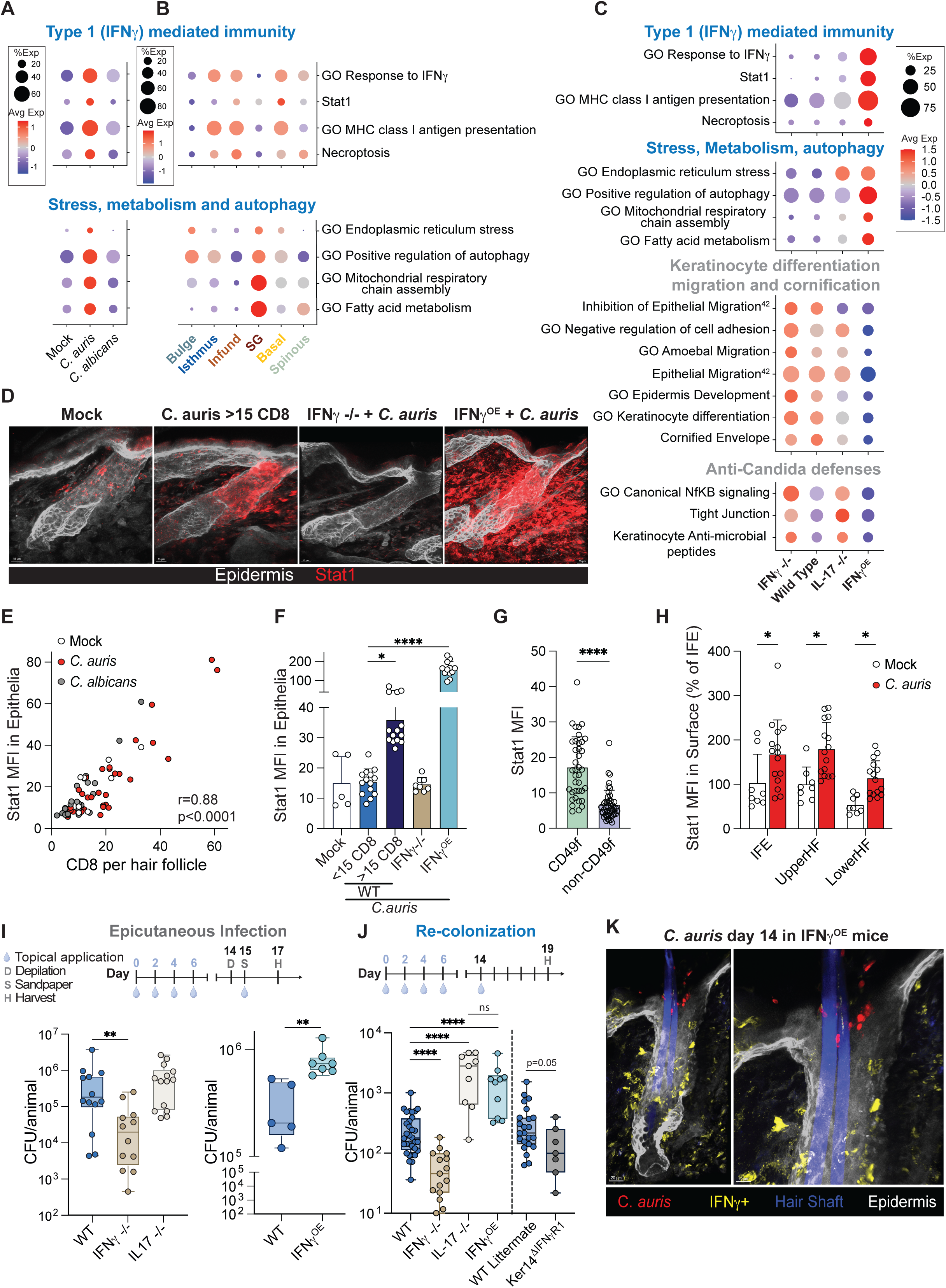
*C. auris* induces IFNγ-mediated epithelial rewiring and enhanced fungal colonization. **(A-C)** scRNAseq dot plot of epithelial cells separated by **(A)** fungal colonization type or **(B)** epithelial cell type or **(C)** IFNγ^-/-^, WT, IL-17^-/-^, and IFNγ^OE^ genotype. Mice were treated with *Candida spp* indicated on days 0,2,4,6 and back skin was harvested on day 14. **(D)** Representative 3D confocal microscopy of Stat1 expression in the upper hair follicle in WT mice treated with either mock colonization or *C. auris* colonization on day 14. **(E)** Correlation between Stat1 expression, assessed via mean fluorescence index (MFI), and CD8 T cells per hair follicle. **(F)** Quantitation of Stat1 mean fluorescence intensity (MFI) per hair follicle in mice indicated. WT *C. auris* colonized mice were subdivided based on having less than or greater than 15 CD8 T-cells per hair follicle. **(G)** Stat1 MFI in the epithelial surface (CD49f) compared to all other skin structures (non-CD49f). **(H)** Stat1 MFI between the interfollicular epithelium (IFE), upper hair follicle, and lower hair follicle in mock colonized (white) and C. auris colonized (red) mice. **(I,J)** Schematics showing models used to test colonization and epicutaneous infection. For epicutaneous infection, intact back skin was pre-colonized with *C. auris* on day 0,2,4,6; on day 14, hair was removed, and the skin was abraded with sandpaper. *C. auris* was reapplied on day 15 and back skin was harvested 2 days later for fungal burden. For ‘Re-Colonization’, intact back skin was pre-colonized with *C. auris* on days 0,2,4,6 and intact back skin was re-colonized on day 14. Back skin was harvested for fungal CFU 5 days after final colonization. Graph showing fungal burden (CFU) after *C. auris* **(I)** epicutaneous infection and **(J)** recolonization of B6 wild type (WT), IFNγ^-/-^, IL-17^-/-^ mice, IFNγ^OE^ mice (Yeti/+), and Ker14^ΔIFNγR1^(Ker14^Cre^;IFNγR1^Flox^). **(K)** IFNγ overexpressing mice (IFNγ^OE^) colonized with *C. auris*, at day 14 with numerous yeast forms at the hair follicle orifice. *P<0.05, ****P<0.0001 for one-way analysis of variance (ANOVA) and Tukey’s multiple comparison test **(F)**. In (E), statistics shown comparing C. auris <15 CD8 per hair follicle to all other groups. **P<0.01, ***P<0.001 for Kruskal-Wallis test with Dunn’s multiple comparison test **(I,J)**. *P<0.05, **P<0.01 student T-Test **(G,H).** *P<0.05, **P<0.01 Mann-Whitney Test **(I,J)**.

The scRNAseq data suggest that *C. auris-*associated transcriptional responses are concentrated in keratinocytes of the upper hair follicle (**Fig4B, Fig4s1E**). To validate these findings, we assayed the expression of key IFNγ response proteins using confocal microscopy. Compared to mock-colonized animals, Stat1 protein was substantially more abundant within upper hair follicle keratinocytes of WT animals following 14 days of colonization with *C. auris* (**Fig4D**). By comparison, Stat1 levels were sharply reduced in *C. auris-*colonized IFNγ^-/-^ animals and massively upregulated in *C. auris-*colonized IFNγ^OE^ animals. Of note, in both colonized and uncolonized animals, expression of Stat1 protein was largely restricted to hair follicles associated with >15 CD8^+^ T cells, and the level of expression at individual hair follicles correlated directly with the number of associated CD8^+^ cells (**Fig4E,F, Fig4s2A**). Stat1 protein signal was also higher in the epithelia compared to the dermis, and highest in the upper hair follicle and interfollicular epidermis (**Fig4G,H**). The MHC-I component B2 Microglobulin was also upregulated in upper hair follicle keratinocytes following colonization with *C. auris* compared to *C. albicans-*colonized mice (**Fig4s2B-D**). Together, these data suggest that *C. auris* not only activates cDC1, Tc1, and Th1 abutting hair follicles, but also induces local IFNγ-dependent transcriptional changes in hair follicle keratinocytes, including the upregulation of pathways associated with Stat1 signaling, antigen presentation, and keratinocyte stress, as well as decreases in barrier repair and anti-*Candida* immunity.

Next, we evaluated the functional consequences of *C. auris-*induced type 1 or type 3/17 immunity on subsequent fungal colonization. Four topical inocula of *C. auris* were administered over six days to provoke adaptive immune responses in WT, IFNγ^-/-^, IL-17^-/-^, and IFNγ^OE^ mice, followed by administration of a rechallenge dose on day 14 or 15 and quantitation of subsequent fungal skin titers (**Fig4I-J**). In separate experiments, rechallenge was performed using two different protocols: (1) epicutaneous infection, in which the inoculum was applied to skin damaged by chemical depilation and sandpaper abrasion to foster higher fungal titers, and (2) standard colonization, in which the inoculum was applied to undamaged skin. We observed no change in *C. auris* skin titers in IL-17^-/-^ animals relative to WT animals following rechallenge using an epicutaneous infection protocol (**Fig4I**); however, skin titers were ∼10-fold higher in the IL-17^-/-^ animals when rechallenge was performed on undamaged skin (**Fig4J**). By contrast, *C. auris* skin titers were significantly reduced in IFNγ^-/-^ mice and significantly elevated in IFNγ^OE^ mice regardless of the rechallenge protocol (**Fig4I-K**), suggesting that IFNγ paradoxically promotes the growth of *C. auris* on skin. Consistent with the findings that IL-17 inhibits but IFNγ promotes *C. auris* skin colonization, we observed that *C. auris* strains that induce stronger IL-17 responses relative to IFNγ exhibit lower skin titers in an epicutaneous infection model (**Fig4s3A,B**).

In aggregate, our findings suggest that IL-17 signaling triggers sterilizing immune responses against *C. albicans* and, to some degree, *C. auris,* whereas IFNγ signaling selectively promotes the growth of *C. auris* in the skin. To clarify whether IFNγ directly benefits *C. auris* or does so indirectly through suppression of IL-17 signaling, we assessed *C. auris* skin titers in colonized animals from which one or both cytokines had been depleted. WT and IL-17^-/-^ mice were treated with IFNγ-blocking antibodies and four topical doses of *C. auris* to undamaged skin, followed by a single topical rechallenge dose to undamaged skin on day 14 and quantification of fungal skin titers on day 19. Treatment with IFNγ-blocking antibodies led to reduced *C. auris* CFU in both WT and IL-17^-/-^ mice (**Fig4s3C)**, suggesting that IFNγ promotes the fitness of *C. auris* in skin independently of IL-17, at least in part. In contrast to skin colonization, during systemic fungal infection with *C. albicans*^37,38^ or *C. auris* (**Fig4s3D**), IFNγ and IL-17 both contribute to protection.

Nonetheless, our scRNAseq data indicate that *C. auris* and *C. albicans* trigger opposing transcriptional responses in hair follicle keratinocytes, with *C. auris* triggering suppression and *C. albicans* activation of genes encoding antifungal defense and barrier integrity factors. Our findings show that IL-17 signaling is directly responsible for the activation of *C. albicans* response genes in keratinocytes. To test whether IFNγ also acts directly on keratinocytes to promote *C. auris* persistence, we generated mice that lack the IFNγ receptor specifically in keratinocytes (Ker14^Cre^; IFNγR1^F/F^ or Ker14^ΔIFNγR1^). Following colonization with *C. auris,* skin titers were significantly reduced in Ker14^ΔIFNγR1^ mice compared to littermate control animals (**Fig4J**), indicating that *C. auris* skin colonization is supported by induced IFNγ signaling that acts directly on skin keratinocytes.

## Discussion

Since its identification in 2009, *C. auris* has appeared on ‘watch lists’ issued by the World Health Organization and US Centers for Disease Control and Prevention of the most worrisome microbial pathogens. *C. auris* exhibits a combination of severe antibiotic resistance and facile person-to-person transmission that is unprecedented among fungal pathogens and more closely resembles the behavior of certain highly transmissible, multidrug-resistant bacterial pathogens. Although investigations of several *C. auris* outbreaks have identified skin colonization as a key factor promoting both pathogen transmission and risk for fungal sepsis, there are no effective procedures to decolonize skin. Meanwhile, basic research on *C. auris* skin colonization remains in its infancy, and two fungal adhesins (Scf1 and Iff4109) are the only molecules known to facilitate the interaction of *C. auris* with mammalian skin^39^.

Against this backdrop, our discovery that *C. auris* induces type 1 immunity in skin and that this immune response paradoxically supports persistence of the fungus was unexpected, particularly as it differs from previous observations with *C. albicans.* Using *C. albicans* for comparison, we reproduced previous reports that *C. albicans* induces sterilizing type 3/17 immune responses in skin. Further, using Ker14^ΔIL17Ra^ mice, we established that IL-17 signaling in keratinocytes is critical to preventing *C. albicans* overgrowth and severe skin pathology in our mouse model and that *C. albicans*-driven IL-17 stimulates the upregulation of antifungal defense genes and genes supporting barrier function in upper hair follicle keratinocytes. By contrast, our analysis of diverse *C. auris* isolates exposed a species-specific hair-binding activity and a strong tropism to hair follicles, with *C. auris* colonizing ∼30% of hair follicles in undamaged skin. Using thick section 3D confocal microscopy, we also mapped the topography of skin immune cells, observing that *C. auris* triggers the proliferation of cDC1, Tc1, Th1 in close approximation to the upper region of colonized hair follicles. Finally, we found that *C. auris* and downstream IFNγ provoke the downregulation of antifungal defense and barrier integrity genes in upper hair follicle keratinocytes—virtually opposite to the response elicited by *C. albicans* and IL-17—and that IFNγ signaling in keratinocytes enhances the ability of *C. auris* to colonize skin. Based on these results, we propose that *C. auris* enhances its own fitness in skin by inducing a localized type 1 immune response to create a hospitable niche for persistence within hair follicles.

Over millennia of coevolution, fungi have developed diverse strategies to thrive in natural environments and to colonize or infect diverse hosts. Still, the interactions of *C. auris* and *C. albicans* with mammalian skin are strikingly different, particularly for species related closely enough to share the same noncanonical genetic code^40,41^. *C. albicans* is a highly adapted commensal of humans and other mammals that primarily inhabits the gut, and invasion into deeper tissues is dramatically restricted by IL-17-dependent immunity. In contrast, although *C. auris* is new to humans, it exhibits a remarkable facility for propagation and persistence on human skin. Our observations suggest *C. auris* is far less sensitive to IL-17-dependent restriction on skin than *C. albicans*. Moreover, it displays unique abilities to bind hair, inhabit hair follicles, and induce highly focal IFNγ-dependent immune responses that reduce antifungal defenses and induce epithelial stress within keratinocytes of the upper hair follicle. Although the variables driving the recent emergence of *C. auris* into human populations remain mysterious, the multiplicity of mechanisms supporting the success of *C. auris* on the skin is suggestive of previous selection in a non-human mammalian host.

## Acknowledgments

We thank Dr. Hiten Madhani and Dr. Richard Locksley for their helpful comments and suggestions.

## Funding

Dermatology Foundation Career Development Award (EDM)

Dermatology Foundation Dermatology Investigator Research Fellowship (EDM)

NIH grant T32AR007175-44 (EDM)

NIH grant T32AR079068 (CH)

National Institute of Neurological Disorders and Stroke grant R01NS126765 (ABM)

UCSF Department of Laboratory Medicine Discretionary Funds (ABM)

National Institute of Allergy and Infectious Diseases grant R01AI180438 (ABM)

National Institute of Arthritis and Musculoskeletal and Skin Diseases grant R01AR084863 (SN/ABM)

National Institute of Allergy and Infectious Diseases grant R21AI171789 (SMN)

## Author contributions

Conceptualization: EDM, VP, SMN, ABM, MDR, TCS

Methodology: EDM, VP, SMN, ABM

Investigation: EDM, VP, PM, AR, CH, EW, PB

Data curation: EDM, VP

Funding acquisition: EDM, SMN, ABM

Writing: EDM, VP, SMN, ABM

Supervision: SMN, ABM

## Competing interests

The authors declare no competing interests.

## Data and materials availability

Single cell transcriptomic data generated in this paper are deposited in Gene Expression Omnibus (GEO) (accession number will be released at the time of publication).

## Supplementary Materials

Materials and Methods

Figs. 1S1-1S2, 2S1-2S3, 3S1, 4S1-4S3

References (1-43)

Movies S1

## FIGURE LEGENDS

**Figure 1S1.**
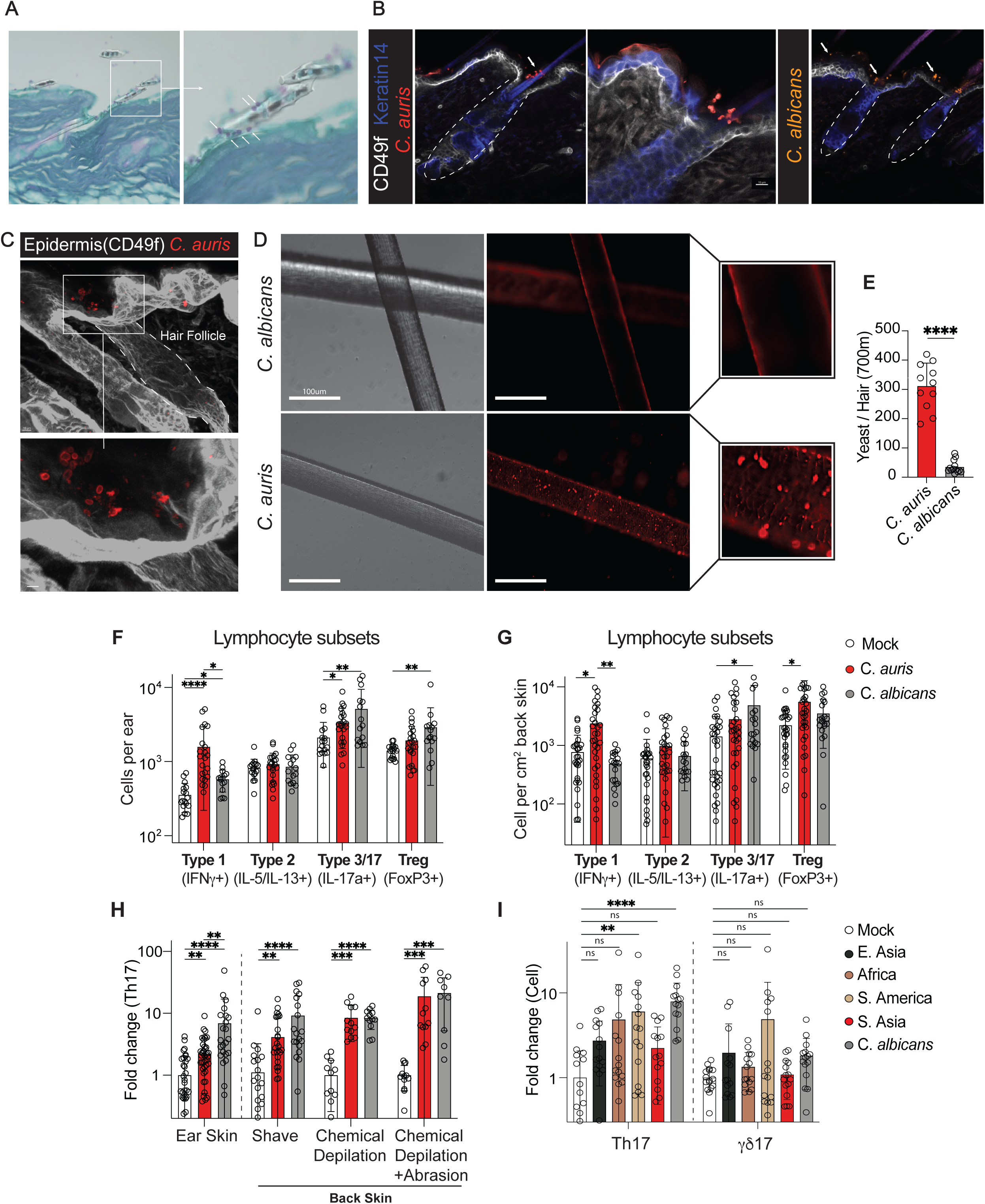
**(A)** PAS fungal staining 2 days after a single inoculum of *C. auris* to intact back skin showing yeast adhering to the hair shaft. **(B)** Virtual slice of a thick section confocal microscopy image of skin with hair follicles, showing *C. auris* or *C. albicans* 2 days after single inoculum of yeast on intact back skin. **(C)** 3D volumetric microscopy image of *C. auris* colonizing the hair follicle orifice at day 14 in repetitive colonization model (63x objective). **(D,E)** Human hair incubated in vitro with either *C. auris* or *C. albicans*, stained with calcofluor white, washed and imaged via fluorescence microscopy (40x), and quantified. (**F,G**) Flow cytometric quantification of lymphocytes (CD45^+^CD90.2^+^) expressing either IFNγ, IL-5 or IL-13, IL-17a, or FoxP3 in the **(F)** ear skin or **(G)** back skin following *C. auris* or *C. albicans* colonization. The fold change is normalized to the same experiment mock-treated controls. **(H)** Flow cytometric quantification in mice treated with various levels of epithelial damage (intact ear skin, intact shaved back skin, back skin depilated with Nair^TM^, back skin depilated with Nair^TM^, and abraded with sandpaper (SP). **(I)** Flow cytometric quantification of Th17 and γδ T cells expressing IL-17 (γδ17) as a fold change compared to same experiment controls across several strains from different clades of *C. auris*. *P<0.05, **P<0.01, ***P<0.001, ****P<0.0001 in student’s t-test **(E)**, one-way analysis of variance (ANOVA) and Tukey’s multiple comparison test **(F,G,H)**, and Kruskal-Wallis test with Dunn’s multiple comparison test **(I)**. All immunophenotyping used a repetitive colonization model with yeast application on day 0,2,4,6 and tissue harvest on day 14.

**Figure 1S2.**
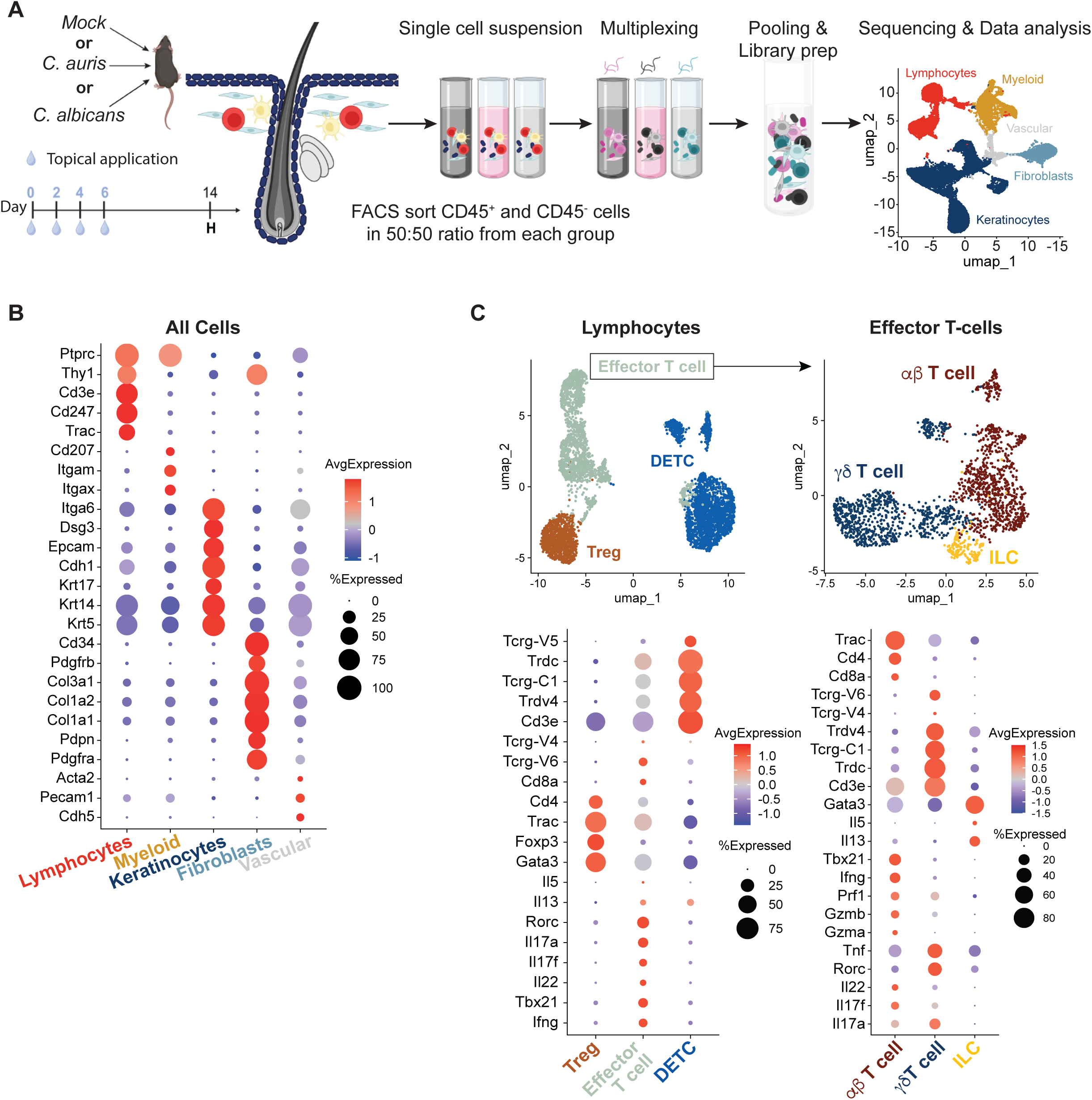
**(A)** Schematic of the workflow for single-cell RNA sequencing (scRNAseq). The back skin of mice was colonized with *C. auris*, *C. albicans,* or PBS controls on day 0,2,4,6; tissue was harvested on day 14, creating a single-cell suspension. Two mice were pooled for each group prior to sorting. Hematopoietic (CD45^+^) and non-hematopoietic (CD45^-^ live cells) were sorted in a 1:1 ratio and pooled together for each sample. Samples were incubated with oligonucleotide barcodes, pooled in equal ratios into one sample for library preparation. Completed libraries were sequenced, aligned with CellRanger multi and analyzed using Seurat 5.0.2. **(B)** Heatmap dot plot of lineage-defining markers for lymphocytes, myeloid cells, keratinocytes, and fibroblasts. **(C)** Strategy for sub-setting T cells based on scRNAseq lineage-defining genes.

**Figure 2s1.**
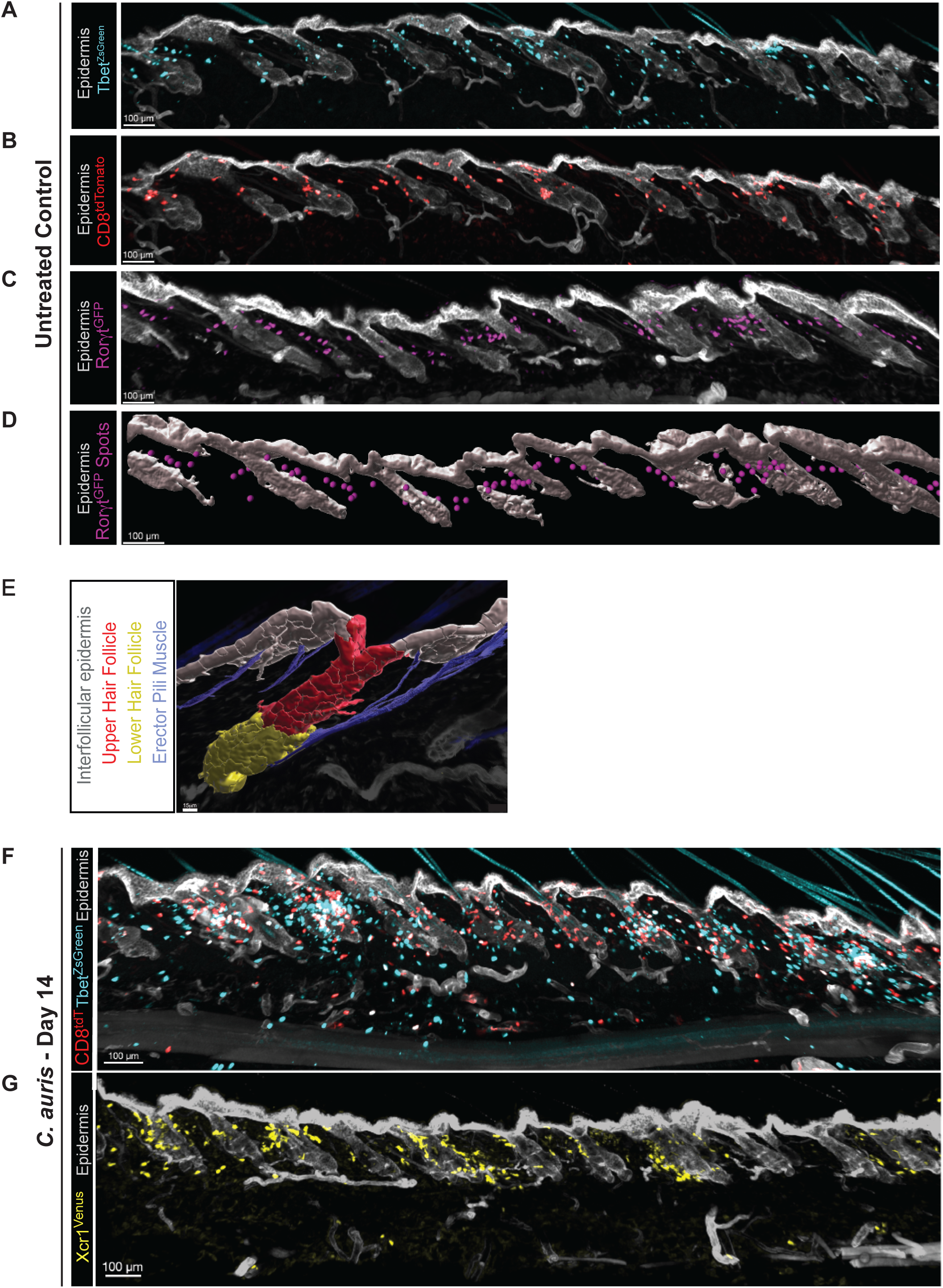
(A-C) Confocal microscopy thick-section imaging of back skin in telogen showing (**A**) distribution of total type 1 lymphocytes (Tbet^ZsGreen^), **(B)** Cytotoxic CD8^+^ T cells (CD8^Cre^;R26^TdTomato+^), or **(C)** type 3/17 lymphocytes (Rorgt^GFP^). **(D)** Representative surfacing on T cell subsets and CD49f epithelia via Imaris software. **(E)** Surfacing of hair follicle niches, derived from native stains and morphologic features. The epidermis (CD49f) was subdivided into interfollicular epidermis (InterEPI), upper hair follicle, lower hair follicle. The epidermis is defined using CD49f (ITGA6), the nerve is defined using PGP9.5 neuronal stain, and the Erector pili muscle is defined using αSMA staining and morphology. **(F,G)** Confocal microscopy thick-section images 14 days post C. auris colonization, using (**E**) CD8^Cre^;R26^TdT+^ mice crossed with Tbet^ZsGreen^ mice or **(F)** Xcr1^Venus^ mice.

**Figure 2s2.**
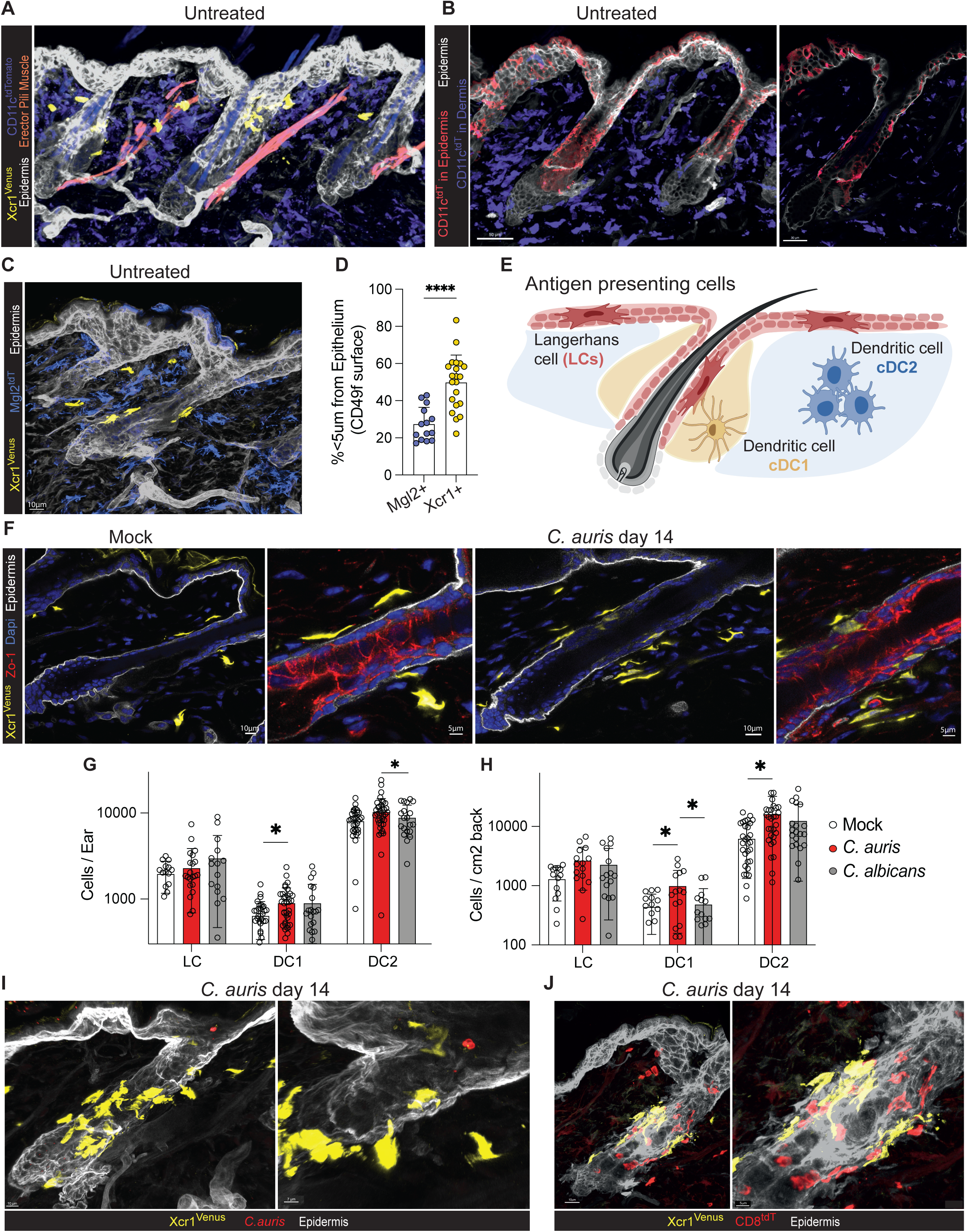
**(A,B,C,D)** Representative confocal microscopy thick section image of hair follicle from untreated **(A,B)** CD11c^Cre^;R26^tdT^; Xcr1^Venus^ mice, (**C**) Mgl2^Cre^;R26^tdT^; Xcr1^Venus^ mice (60x) with **(D)** quantitation of cells that reside <5um from CD49f epithelial surface. Each dot represents an image from a single hair follicle. **(B)** Showing virtual thin-section highlighting both epithelial (Langerhans cells) and dermal-associated CD11c^+^ cells. **(E)** Schematic of proposed topography of Antigen Presenting Cells (APC) subsets in skin. **(F)** Representative confocal microscopy thick section image of mock-treated versus *C. auris* treated back skin with cDC1 (Xcr1^Venus^) surrounding the upper hair follicle and with projections protruding between the keratinocytes near tight junction protein ZO-1. **(G,H)** Antigen-presenting cells (APC) were quantified via flow cytometry in **(G)** ear or **(H)** back skin. **(I-J)** Representative confocal microscopy thick section image of *C. auris* **(I)** within the hair follicle orifice near cDC1 and **(J)** cDC1 colocalizing with CD8^+^ T cells (CD8^Cre^;R26^TdT^;Xcr1^Venus^) at day 14. ****P<0.0001 student T-Test **(B).** Images are representative of 3 independent experiments.

**Figure 2s3.**
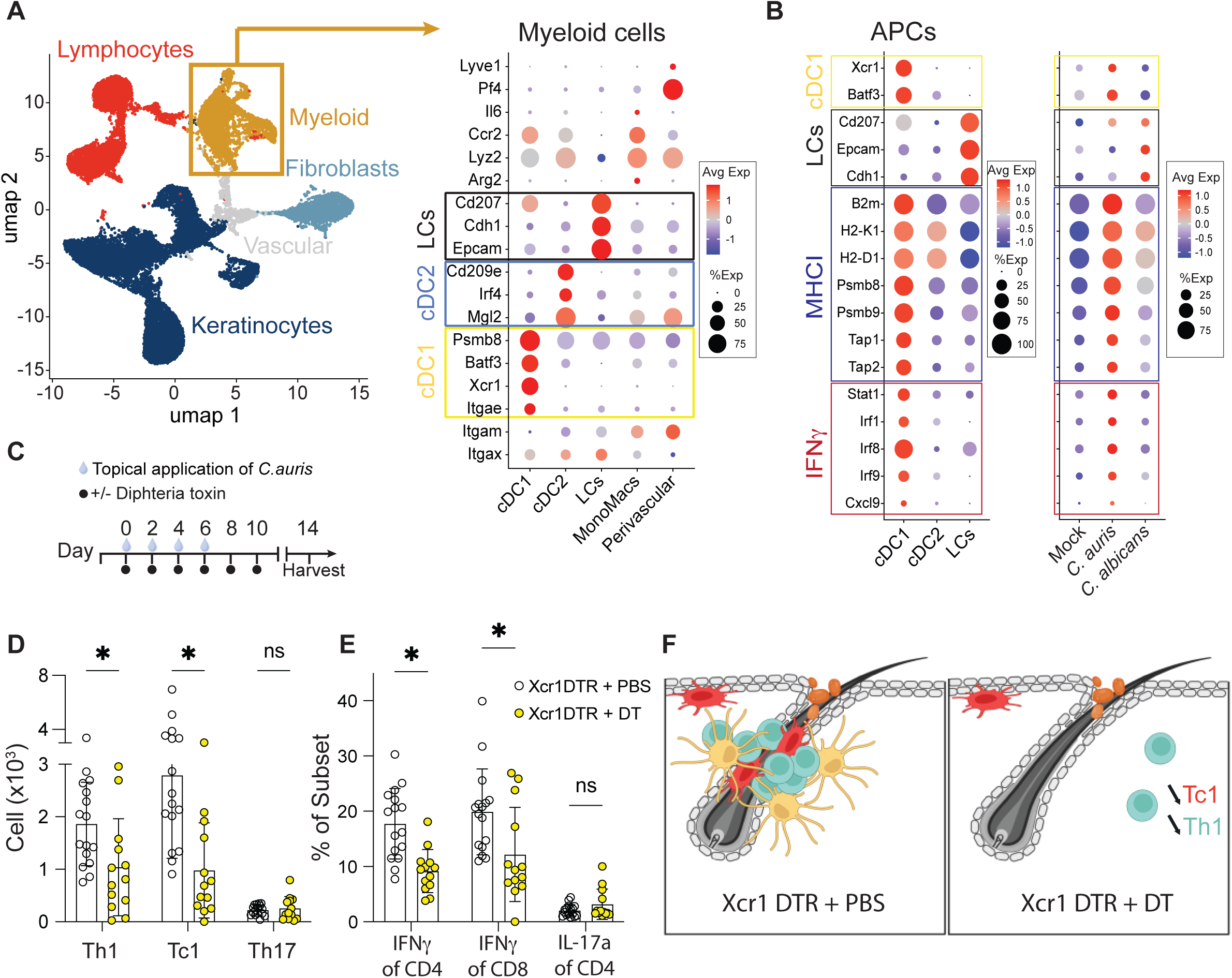
**(A)** From scRNAseq of mock-treated, *C. auris,* or *C. albicans* colonized mice (repetitive colonization on day 0,2,4,6 and harvest on day 14), a large population of ‘myeloid’ cells (CD45^+^ cells lacking T-cell markers) were divided into subsets based on defined transcriptional signatures. **(B)** Pre-selected on all APC (CD45^+^CD11c^+^MHCII^+^ cells) and subdivided as Langerhans cells (LC, EpCAM^high^), cDC1 (EpCAM^-^CD11b^low^CD103^high^) and cDC2 (EpCAM^-^CD11b^var^CD103^low^). Dot plot showing relative gene expression of selected genes in APC subsets (left panel) and in mock, C. auris or C. albicans colonized mice (right panel). **(C-D)** Xcr1^DTR^ mice were treated with diphtheria toxin (DT) or PBS on days 0,2,4,6,8 and 10 during colonization with *C. auris* and mouse ear skin was harvested at day 14. Flow enumeration of **(C)** number and **(D)** Percent cytokine production of Th1 (CD4^+^IFNγ^+^ T cells), Tc1 (CD8^+^IFNγ^+^ T cells), and Th17 (CD4^+^IL-17a^+^ T cells) in mice colonized with *C. auris* and treated with DT or PBS. **(E)** Cartoon schematic depicting loss of cDC1 leading to decreased type 1 upper hair follicle centered immune response. *P<0.05, **P<0.01, ***P<0.001, ****P<0.0001 for one-way analysis of variance (ANOVA) and Tukey’s multiple comparison test **(A,B)**. *P<0.05, student T-Test **(F,G).**

**Figure 3s1.**
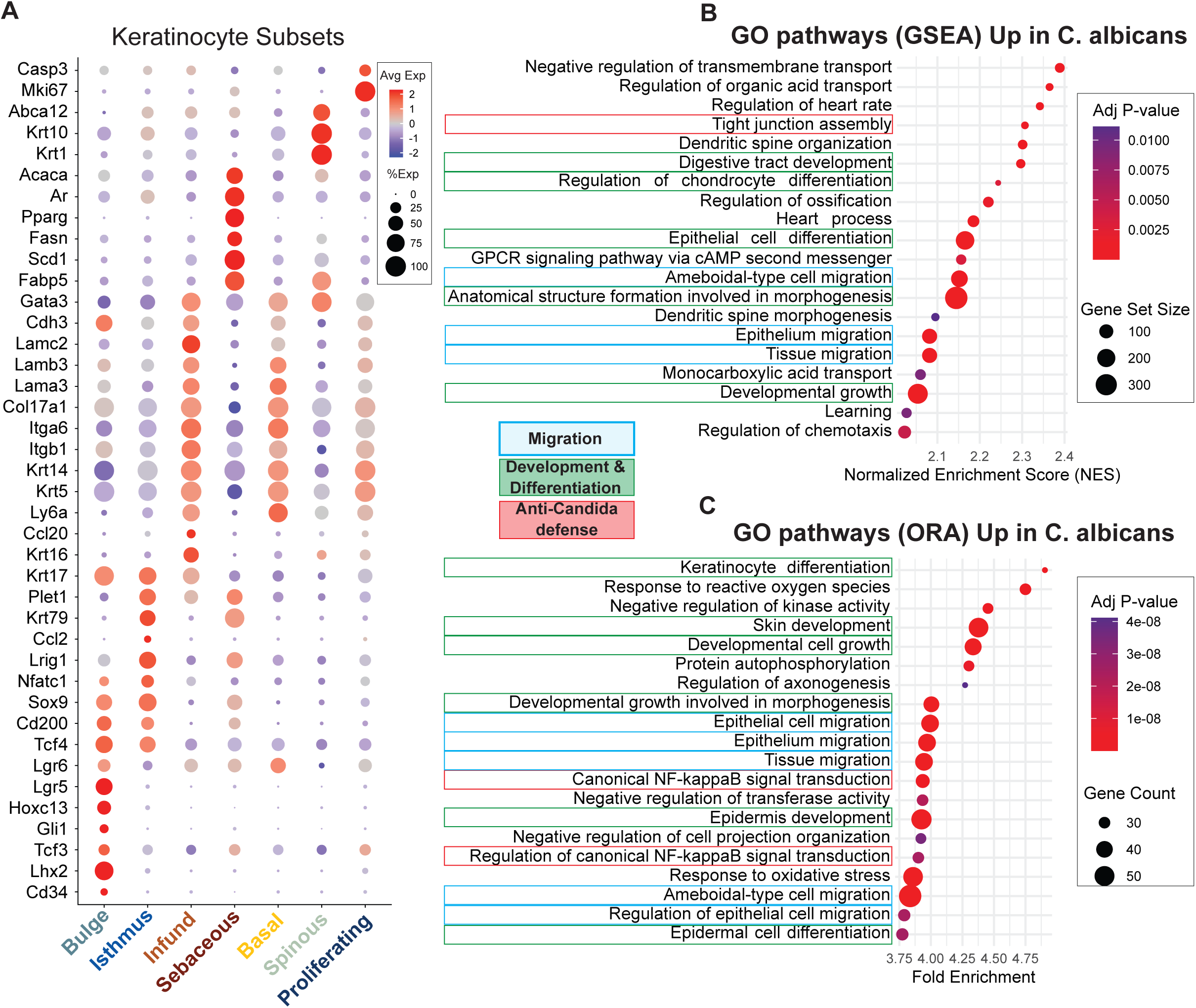
**(A)** scRNAseq dot plot of keratinocyte genes used to define keratinocyte subsets: bulge, isthmus, infundibulum, basal layer, spinous layer, and sebaceous gland. **(B)** Gene Set Enrichment Analysis (GSEA) showing upregulated GO terms after *C. albicans* colonization. **(C)** Over-Representation Analysis (ORA) showing upregulated gene ontology (GO) terms after *C. albicans* colonization.

**Figure 4s1.**
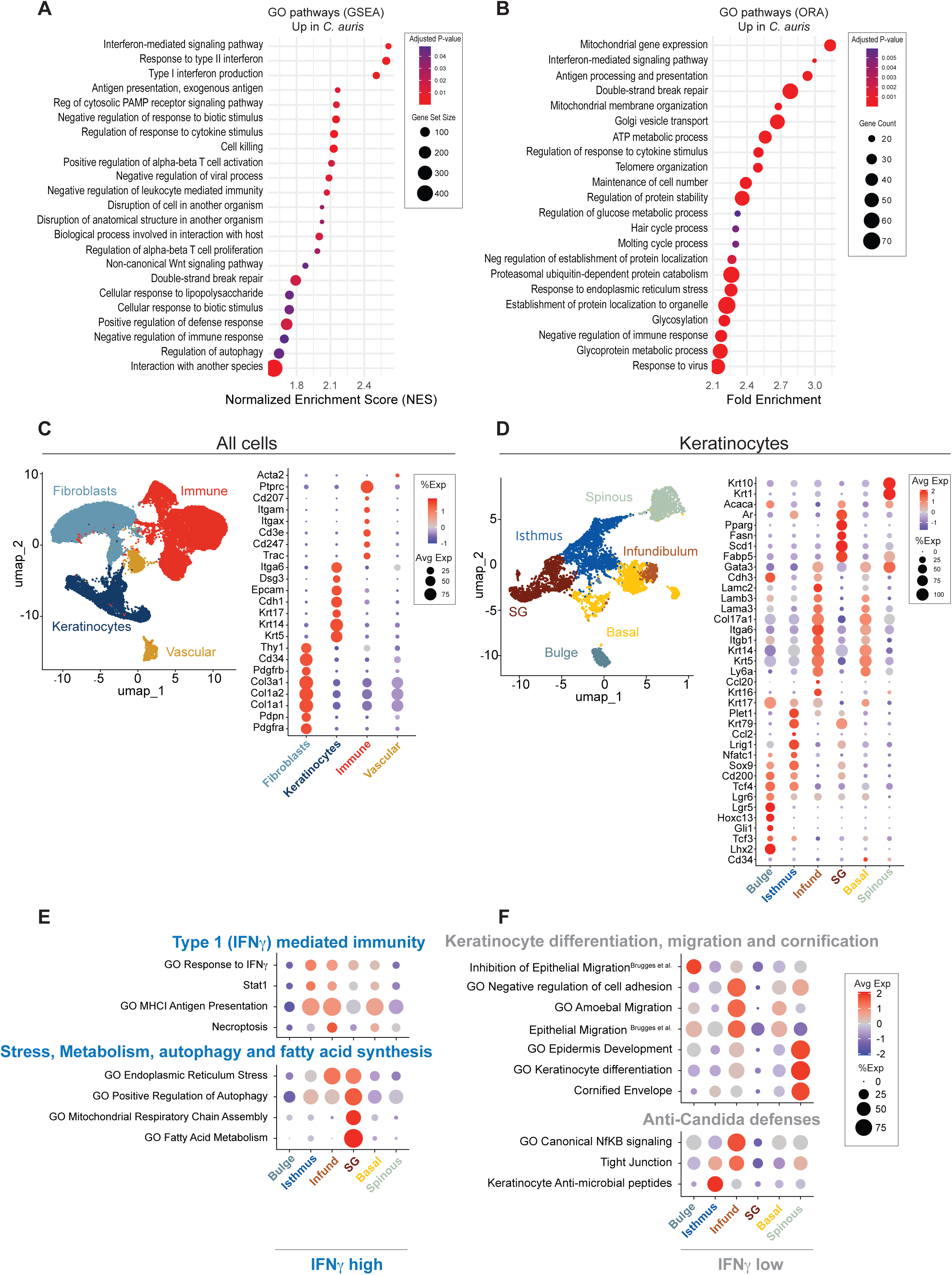
**(A,B) (A)** Gene Set Enrichment Analysis (GSEA) of scRNAseq showing GO terms after *C. auris* colonization or **(B)** Over-Representation Analysis (ORA) showing upregulated gene ontology (GO) terms after *C. albicans* colonization. **(C)** scRNAseq Umap (left panel) showing annotated clusters revealing large population of keratinocytes, fibroblasts, immune cells and vascular cells. Dot plot (right panel) showing lineage defining markers each cell type. **(D)** scRNAseq Umap (left panel) gated on keratinocytes showing keratinocyte subsets and dotplot (right panel) of putative keratinocyte gene makers used to define keratinocyte subsets: bulge, isthmus, infundibulum, basal layer, spinous layer and sebaceous gland. (**E,F**) scRNAseq dotplot by cell type showing modules upregulated in **(E)** IFNγ^OE^ or **(F)** IFNγ -/- mice.

**Figure 4s2.**
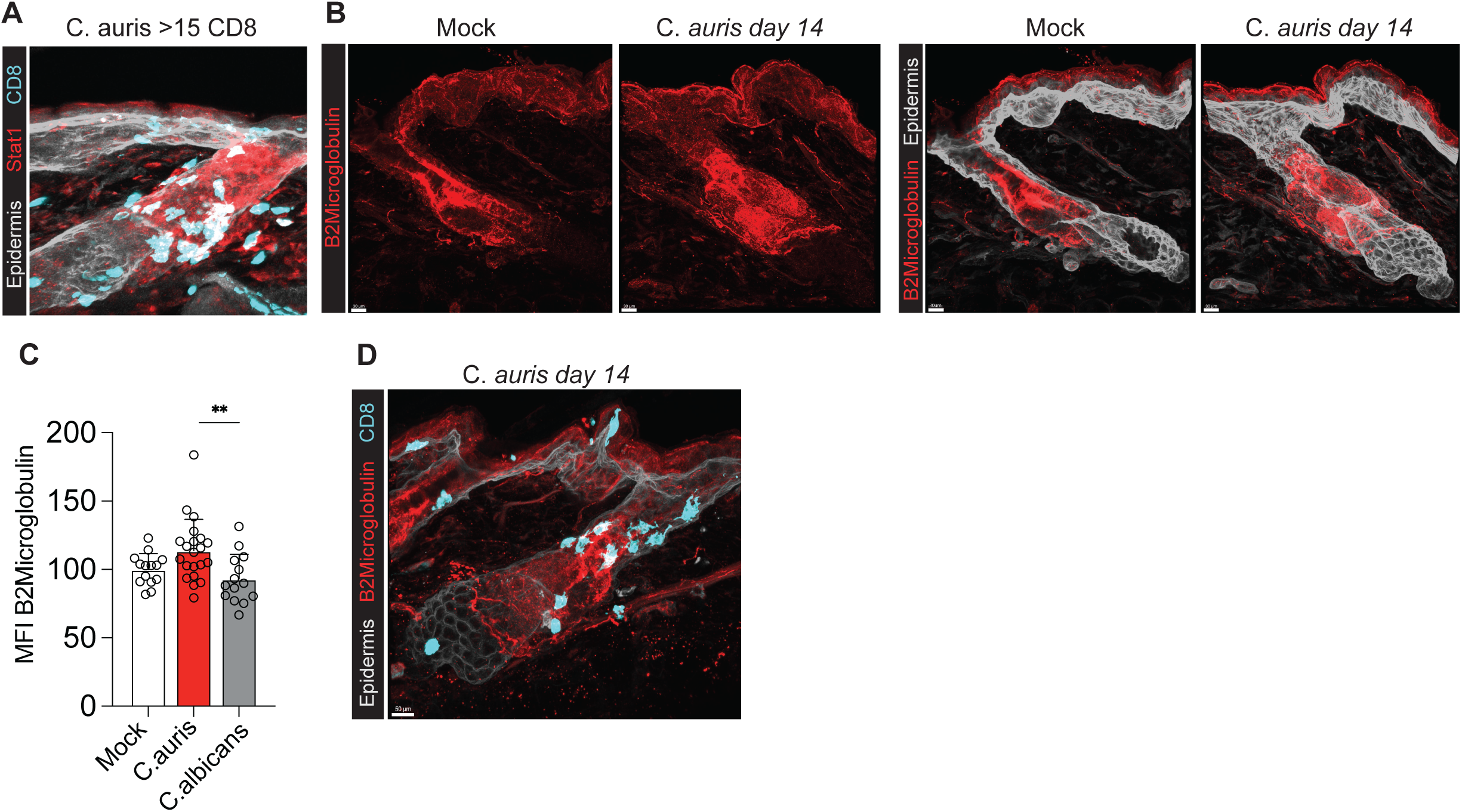
**(A)** Representative confocal microscopy thick-section image of the overlay of CD8^+^ T cells (CD8^Cre^;R26^TdT+^) and Stat1 staining in a hair follicle with >15 CD8^+^ T cells after *C. auris* colonization. **(B)** Representative thick-section image showing Beta2 Microglobulin in the upper hair follicle of WT mice comparing mock colonized to *C. auris* colonized mice on day 14. **(C)** Image quantitation of Beta2 Microglobulin MFI in epithelial surface comparing mock-treated, *C. auris* colonized, and *C. albicans* colonized mice on day 14. **(D)** Representative thick-section image of overlay of CD8^+^ T cells and Beta2 Microglobulin staining. **P<0.01 for one-way analysis of variance (ANOVA) and Tukey’s multiple comparison test **(C,D)**.

**Figure 4s3.**
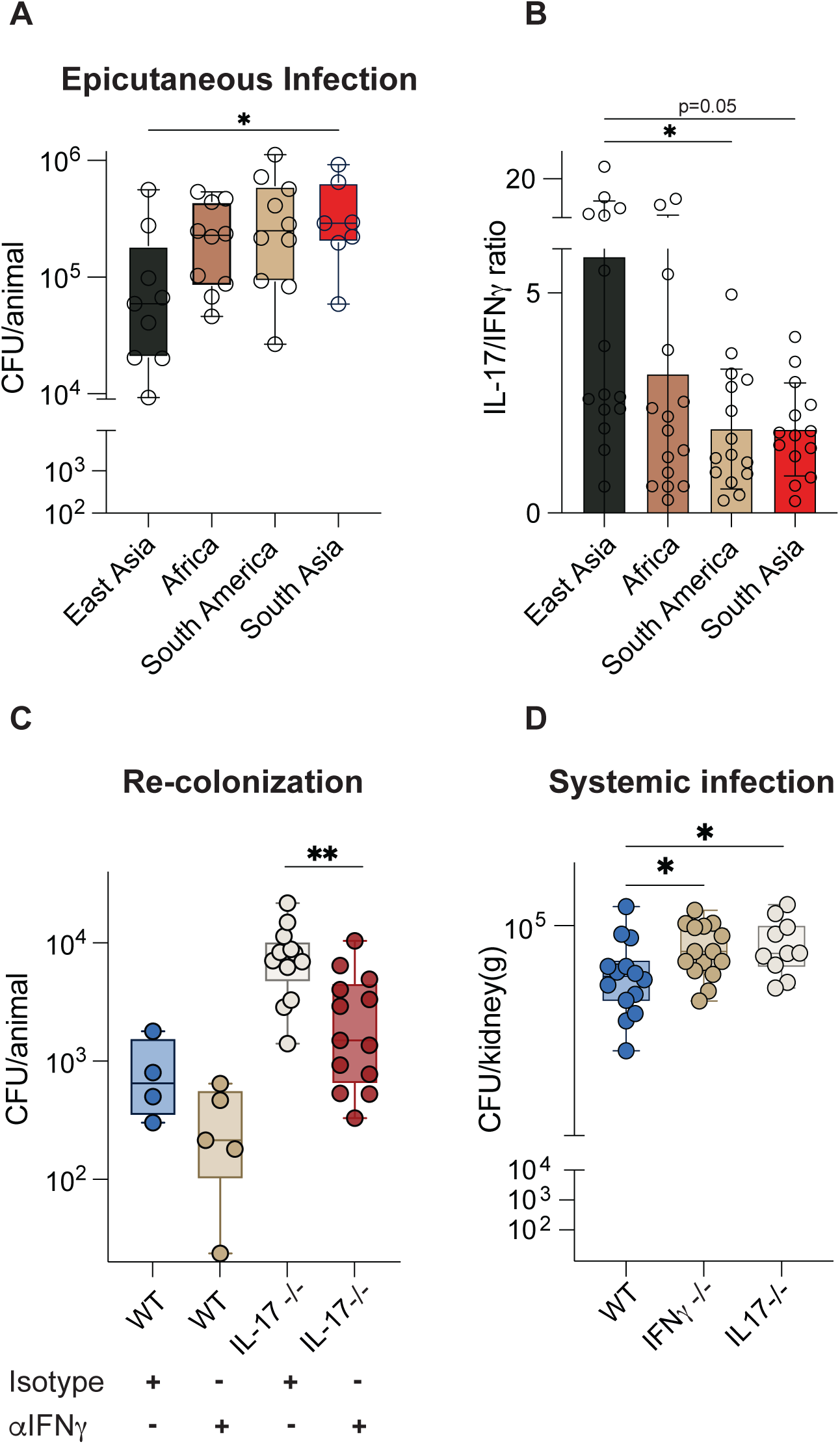
**(A)** Epicutaneous rechallenge model (Fig4A) showing CFU of different clades of *C. auris*. **(B)** Flow cytometric quantification of IL-17a to IFNγ lymphocyte ratio in ear skin of mice colonized by different clades of *C. auris*. **(C)** Wild type or IL-17 -/- mice were treated with either anti-IFNγ therapy or isotype control throughout colonization with *C. auris* on days 0,2,4,6 and recolonization on day 14. Back skin was harvested for fungal burden 5 days following recolonization. **(D)** Mice were injected with 5×10^7^ CFU retro-orbitally of *C. auris* or C. albicans and kidney was harvested 48 hours later and CFU determined in B6 wild type (WT), IFNγ^-/-^, or IL-17^-/-^ mice. *P<0.05, **P<0.01 for the Kruskal-Wallis test with Dunn’s multiple comparison test **(A,B,D)**. For **(A,B)** South Asia was compared to all other strains. **P<0.01 Mann-Whitney Test **(C).** For **(D)** IFNγ and IL-17-/- were compared to WT controls.

**Supplemental Movie 1** Thick-section confocal image of *C. auris* colonized skin at day 14 showing native stains and Imaris generated surfaces and spots. Epidermis (CD49f, white), erector pili muscle (aSMA, blue), hair shaft (autofluorescence, orange), sebaceous gland (Fabp5, yellow), *C. auris* (red), and Tbet+ Lymphocytes (Tbet^Zsgreen^, cyan).

## METHODS

### Mice

C57BL/6 mice from Jackson Laboratories mice were used for immunophenotyping experiments. We used mice lacking IFNγ (IFNγ^-/-^) from Jackson (B6.129S7-Ifngtm1Ts/J; MGI:J:66802), mice overexpressing IFNγ (IFNγ^OE^) generously gifted from Richard Locksley lab (Mouse: B6.129S4-Ifngtm3.1Lky/J (YFP-Enhanced Transcript for IFN-γ or Yeti), mice lacking IL-17a and IL-17f (IL-17af^-/-^) from Jackson (B6.Cg-Il17a/Il17ftm1.1Impr Thy1a/J, MGI:J:187400). To generate keratinocyte specific knock out of IL-17 and IFNγ signaling, we crossed Ker14^Cre^ mice (B6N.Cg-Tg(KRT14-cre)1Amc/J) from Jackson with Il17Ra floxed mice (B6.Cg-Il17ratm2.1Koll/J; MGI:J:237978) and IFNγR1 floxed mice (C57BL/6N-Ifngr1tm1.1Rds/J, MGI:J:206257) from Jackson Labs.

To track type 1 lymphocytes we used Tbet (Tbx21)-zsGreen transgenic mice (generously provided by Jinfang Zhu, Lab of Immune System Biology, NIH). To track type 3/17 lymphocytes we used Rorc(gamma-t) eGFP mice (Rorgt^GFP^) (generously provided by Gerard Eberl; MGI 3829387). To track CD8+ lymphocytes, we used CD8^Cre^, which has cre recombinase activity peripheral CD8+ lymphocytes, but not CD4+CD8+ Thymic progenitors, from Jackson (C57BL/6-Tg(Cd8a-cre)1Itan/J, MGI:J:141009). These mice were crossed with Rosa26^TdT-Ai14^ mice (R26-CAG-RFP, Jackson 007914) containing a flox-stop-flox sequence upstream of a CAG-RFP-WPRE cassette in the constitutively expressed ROSA26 locus. To track cDC1s, Xcr1^Venus/+^ mice (Xcr1tm1Ksho; MGI:5544057) were used. To track cDC2s, Mgl2^Cre^ (generously given from Tiffany Scharschmidt lab) and CD11c^Cre^ Jackson (B6.Cg-Tg(Itgax-cre)1-1Reiz/J, MGI:J:125869) were crossed with Rosa26^TdT-Ai14^ mice (R26-CAG-RFP, Jackson 007914) as above.

To deplete cDC1s, we used Xcr1^DTR^ mice from Jackson (Xcr1tm2(HBEGF/Venus)Ksho, MGI:5544058).

For lineage tracing of Lrig1^+^ epithelial cells, we used Lrig1^CreERT^ mice that were crossed with Rosa26^TdT^ mice.

If not otherwise stated, all experiments were performed with 7-12-week-old male and female mice. Older mice were not used to ensure back skin hair follicles were harvested during the telogen hair follicle cycle. All mice were bred and maintained in specific-pathogen-free conditions at the animal facilities of UCSF and were used in accordance with institutional guidelines and under study protocols approved by the UCSF Institutional Animal Care and Use Committee.

### Candida strains

*C. auris*: NIH0388 Clade 1 South Asia, NIH0381 Clade 2 East Asia, NIH0383 Clade 3 Africa, NIH0385 Clade 4 South America. *C. albicans*: SC5314.

### Organism and inoculum preparation for colonization

Strains were streaked from glycerol stock solutions onto YEPD media and incubated for 2 days at 30°C. The plates were kept at room temperature for no more than 7 days*. C. albicans* and *C. auris* cultures were initiated from overnight liquid cultures and diluted to an optical density (OD) of 0.1 at 600 nm in glucose-yeast extract-peptone (YEP) broth (containing 2% glucose, 1% peptone, and 0.3% yeast extract). These cultures were then grown at 30°C until the OD at 600 nm reached between 0.7 and 1. Yeast cells were harvested, washed, enumerated using a hemocytometer, and diluted to a concentration of 5 × 10⁸ cells/mL in saline solution for immediate use. The viable counts of the inocula were confirmed by plating serial dilutions on YPD agar plates.

### Fungal colonization and infection models

The dorsal area of the mice was shaved one day prior to colonization using an electric razor. 200 μL or 70ul of *C. albicans* or *C. auris (*concentration of 5 × 10^8^ cells/mL) was applied to back skin and ear skin, respectively. *Candida* was then gently spread using a cotton tip applicator.

#### CFU time course

Mice were colonized with a single application of yeast on day 0 then tissue was harvested for CFUs on days 2,5,8,10,12 or mice were colonized every other day (days 0,2,4,6) and tissue was harvested CFU on days 8, 10 and 14.

#### Re-colonization model for CFU

After shaving as above, mice were pre-colonized with yeast on days 0,2,4,6. On day 14, mice underwent a rechallenge by directly reapplying the 70ul of yeast to intact back skin. Mice were monitored for an additional five days before euthanasia and tissue harvest of back skin for CFUs.

#### Epicutaneous infection

After shaving as above, mice were pre-colonized with yeast on days 0,2,4,6. On day 14, depilatory cream (Nair^TM^) was applied for one minute to ensure complete hair removal. On day 15, mice were treated with sandpaper abrasion, applied in three gentle passes, followed by the reapplication of 70ul of yeast. Mice were then observed for two additional days before euthanasia and tissue harvest of back skin for CFUs.

#### Systemic infection

A 100 μL of a suspension at a concentration of 5 × 10^8^ CFU/mL was administered via retro-orbital injection. The mice were observed for two days before being euthanized. Both kidneys were harvested, weighed, and homogenized in 1 mL of PBS. The homogenate was then subjected to a series of dilutions and plated onto Sabouraud dextrose agar plates supplemented with gentamycin and ampicillin. Colony-forming units (CFUs) were counted after 2 days of incubation at 30°C.

#### Flow cytometry experiments

For ear skin, 30ul of solution was added to each ear and gently spread using a cotton tip applicator on days 0,2,4,6 and tissue was harvested and processed on day 14. For back skin, one day prior to fungal colonization, skin was either shaved, shaved and depilated, or shaved, depilated and abraded with sandpaper (as above). 200ul of fungus was applied on days 0,2,4,6 and tissue was harvested and processed on day 14.

### Antibody depletion in fungal colonization model

Mice were treated with intraperitoneal injection of 500ug InVivoMAb anti-mouse IFNγ (BioXCell BE0054) or isotype control (BioXCell BE0088) on days 0, 3, 6, 10 and 1g on day 13 (1 day prior to rechallenge).

### Flow cytometry

Resuspended samples were stained in 96-well U-bottom plates. Surface staining was performed at 4°C for 20 minutes in 50μL staining volume. For experiments involving intra-cellular staining, cells were fixed for at 4°C for 20 minutes and permeabilized using BD Cytofix/Cytoperm™ Fixation/Permeabilization Solution Kit for 1 hour. All samples were acquired on a BD LSRII Fortessa Dual or a BD FACSAria II for cell sorting Live cells were gated based on their forward and side scatter followed by Fixable Viability Dye eF780 (eBioScience 65086514) or DAPI (40,6-diamidine-20-phenylindole dihydrochloride, Millipore Sigma D9542-10MG) exclusion. Data were analyzed using FlowJo software (TreeStar, USA) and compiled using Prism (Graphpad Software). Cell counts were performed using flow cytometry counting beads (CountBright Absolute; Life Technologies) per the manufacturer instructions.

Lineages were defined as the following: Dendritic cells were defined as dapi negative, CD45+, CD11c+, MHCII+. Langerhaans cells were EpCAM positive, cDC2s were EpCAM negative, CD103 negative and CD11b low to high. cDC1s were EpCAM negative, CD11b low and CD103^+^. T cells were defined as CD45^+^, CD90.2^+^, CD3^+^. Type 1 lymphocytes were defined as CD45^+^CD90.2^+^IFNγ^+^. Of these, Th1 cells are CD4^+^IFNγ^+^, Tc1 CD8^+^IFNγ^+^. Type3/17 lymphocytes were defined as CD45^+^CD90.2^+^IL-17a^+^. Of these, Th17 were CD4^+^IL-17a^+^, and γδ17 were TCRγδ intermediate IL-17a^+^. Type 2 lymphocytes were CD45^+^CD90.2^+^IL-5^+^ and/or IL-13^+^.

### Flow cytometry antibodies

Antibodies used for flow cytometry include anti CD45 (30-F11, BD Biosciences 564279), anti-CD90.2 (Thy1, 53-2.1, Biolegend 140318, BD Biosciences 553004), anti-CD3ε (17A2, Biolegend 100216), anti-CD4 (RM4-5, Biolegend 100557, or GK1.5, BD Biosciences 563050), anti-CD8α (53-6.7, Biolegend 100750), anti-TCRgamma (GL3, Biolegend 118120), anti-FoxP3 (FJK-16s, eBioscience 53-5773-82), anti-IFNγ (XMG1.2, Biolegend, 505810), anti-IL-17a (TC11-18H10.1, Biolegend, 506922), anti-IL-5 (TRFK5, Biolegend 504304), anti-IL-13 (eBio13A, eBioscience, 12-7133-82), anti-CD11b (M1/70, Biolegend 101224), anti-Ly6G (HK1.4, Biolegend 128005), anti-Ly6C (HK1.4, Biolegend 128011), anti-CCR2 (475301, R&D systems, FAB5538P-100), anti-CD31(390, Biolegend, 102432), anti-CD11c (HL3, BD Biosciences, 558079), anti-MHCII (M5/114.15.2, eBioscience, 56-5321-82), anti-CD103 (2E7, eBioscience, 17-1031-82), CD24 (M1/69, BD Biosciences, 747717), anti-CD64 (X-54-5/7.1, Biolegend 139323), anti-EpCAM (G8.8, Biolegend, 118241), anti-F4/80 (BM8, eBioscience, 67-4801-82), anti-siglecF (E50-2440, BD Bioscience, 740956).

### Colony forming unity enumeration

Back skin was excised, weighed, and homogenized in 3 mL of PBS. The homogenate was serially diluted and plated onto Sabouraud dextrose agar plates containing gentamycin and ampicillin. Plates were incubated at 30°C for 48 hours to quantify CFU as a marker of fungal colonization.

### In vitro hair association

Cultures of *Candida albicans* and *Candida auris* were grown at 30°C until reaching an optical density (OD) of 2. Equal numbers of hair samples of the same size were rinsed with PBS and immersed in the respective cultures, which were supplemented with gentamycin and ampicillin. These cultures were then incubated overnight at 30°C under gentle agitation. The hair samples were retrieved, rinsed in three successive baths of PBS, and stained with Calcofluor White. The stained samples were observed and quantified using microscopy.

### Scanning Electron Microscopy (SEM)

Human hair samples were incubated for 24 hours in YEPD with Candida auris at 30°C. After incubation, the samples were rinsed three times in Phosphate Buffered Saline (PBS) to remove non-adherent yeast cells. The washed hair samples were then affixed to carbon tape on a metal mount. Subsequently, the samples were sputter coated with a 4 nm layer of gold/palladium (Au/Pd). Imaging was conducted using a Sigma VP 500 SEM.

### Tissue processing for flow cytometry

#### Back skin

After euthanasia, back skin was dissected, and the deep fat, panniculosus carnosis muscle, and subcutaneous fat was removed and discarded. 1.5×1.5cm squares were cut, and placed in 12 well plates containing 2ml of digestion media (2 mg ml–1 collagenase XI, 0.5 mg ml–1 hyaluronidase, 0.1 mg ml–1 DNase in RPMI, 1% HEPES buffer, 1% non-essential amino acids, 1% Glutamax and 1% penicillin–streptomycin) finely minced with scissors, digested for 90 minutes at 37 °C. Digestion was then quenched with 3 ml cold RPMI. Samples were strained through 70um filters and a single cell suspension was plated in 96 well round bottom plates.

#### Ear skin

After euthanasia, ear skin was excised and separated into dorsal and ventral sheets and placed dermis side down into 0.5ml digest media (as above) in 24 well plates. Ears were incubated for 60 minutes at 37 °C, finely minced and then incubated for an additional 30 minutes at 37 °C. Digestion was then quenched with 0.5 ml cold RPMI. Samples were strained through 70um filters and a single cell suspension was plated in 96 well round bottom plates.

### Histology

Skins samples were fixed in 10% formalin for 24 hours and washed in 80% ethanol. Preparation of paraffin blocks and 4 μm sections as well as staining with PAS were performed by Nationwide Histology (MT).

### Tissue processing for 3D confocal imaging

Following CO_2_ euthanasia, back skin was removed and floated dermis side down in 2% PFA (Thermo Scientific). After 1 hour (4°C), the skin was submerged under PFA for 6 additional hours (4°C). The skin was then submerged in 30% sucrose overnight. 1cm sqaures were then cut and embedded in O.C.T. (Thermo Scientific) and frozen first with dry ice and then stored at -80°C. Using a cryostat, skin was sliced in 200μm sections were cut, washed in PBS and then blocked (24 hours 37°C, DPBS/0.3% TritonX-100/1% secondary-host serum/1% BSA). Samples were incubated in primary antibodies diluted in blocking solution (37°C, 48-72 hours), washed for 2 hours 3 times and incubated in secondary antibodies (37°C, 24 hours). Sections were washed for 8 hours 3 times and cleared using Ce3D clearing solution (40% n-methylacetamide, and imaged on a Leica SP8 or Nikon A1R confocal microscopes. For quantitative imaging showing proximity to epithelia (Fig 2C), large images were captured at 20x magnification. Each dot represents a single image with ∼10-15 hair follicles and 2 images were taken per mouse. For quantitation of individual hair follicles, higher magnification images (60-63x) were taken of 3-20 hair follicles/per mouse with at least 3 mice per condition.

### Lrig1 lineage tracing

Lrig1^CreERT^;R26^Tdt^ mice were treated with 75mg/kg tamoxifen (Sigma T5648-5G) dissolved in corn oil (Sigma C8267-500ML) on day -1. On day 0 mice were colonized with mock treatment, *C. auris* or *C. albicans* and tissue was harvested for 3D imaging on day 5.

### Imaging Antibodies

Primary antibodies used for murine imaging include chicken anti-GFP (Aves Labs GFP-1020, 1:400), rabbit anti-dsRed (Takara 632496, 1:400), rabbit anti-Venus (Thermo OSE00002W, 1:400), goat anti-RFP (Rockland, 200-101-379, 1:400), rabbit anti-zsGreen (Takara, 632474, 1:400), rat anti-CD49f AF647 (Biolegend, 313610, 1:200), rat anti-CD49f AF488 (Biolegend, 313608, 1:200), rat anti-alpha Smooth Muscle Actin (Biotium, BNC040665, 1:100), rabbit anti-Stat1 (Cell Signaling Technology, 14994T, 1:200), rabbit anti-Zo1 (Invitrogen, 40-2200, 1:200), 9661T, 1:200), rabbit anti-PGP9.5 (Abcam, ab108986, 1:200), rabbit anti-B2Microglobulin (Proteintech, 13511-1-AP, 1:200), goat anti-FABP5 (R&D, AF1476-SP, 1:400), rabbit anti-*Candida albicans* (Abcam, ab252746, 1:200). Secondary antibodies were used as necessary at protocol-specific concentrations, conjugated to A488, A555, AF594, and A647 (Life Technologies, Thermo-Fisher).

### Confocal microscopy

Confocal images (for thick sections and T-cell quantification) were imaged using a Leica SP8 Upright laser-scanning confocal equipped with a 405nm laser and adjustable white light laser for excitation and imaging with 20x water immersion and 63x water immersion objectives. Nikon A1R laser scanning confocal including 405, 488, 561, and 650 laser lines for excitation and imaging with 60X/1.2 NA Plan Apo VC water immersion objectives. For large scans with 20x objective z-steps were taken every 5um and for individual hair follicles z-tiles were taken every 2um.

### Image analysis/quantification

z-stacks were rendered in 3D and quantitatively analyzed using Bitplane Imaris v10.2 software package (Andor Technology PLC, Belfast, N. Ireland). Individual cells (e.g., lymphocytes and dendritic cells) were annotated using the Imaris ‘spots’ function based on fluorescent Tbet^ZsGreen^, Rorgt^GFP^, CD8^Cre^;Rosa26^TdT^, XCR1Venus and Mgl2^Cre^; Rosa26^TdT^ size/morphological characteristics, with background signal in unrelated channels excluded. 3D reconstructions of the epithelia were generated using the Imaris surface function based on fluorescent CD49f staining and morphologic characteristics. The epithelia was subdivided into interfollicular epidermis (IFE), Upper Hair Follicle and Lower Hair follicle based on morphology and aid from insertion of the Erector Pili muscle when available (aSMA staining). 3D distances between immune cells (lymphocytes and dendritic cells) were calculated using Imaris shortest distance to surface from spots feature. Cells were deemed to be within the surface if <0um from surface. Cells were deemed to be adjacent or within the surface if <5um from surface. MFI of individual channels (e.g. Stat1, B2Microglobulin) was calculated within surfaces (mean value was recorded).

### scRNAseq Tissue preparation and cell sorting

#### Experiment 1

After tissue digestion of back skin (as above) and single cell solution formation, CD45+ and CD45-cells were sorted using in a 1:1 ratio from mock colonized B6 mice, *C. auris* colonized B6 mice and *C. albicans* colonized B6 mice (4 applications on days 0,2,4,6 and harvest on day 14). Prior to sorting 2 mice per condition were pooled. After sorting, 2 replicates per group (2x Untreated, 2x *C. auris* colonized, 2x *C. albicans* colonized) were multiplexed in equal ratios using 10x 3’ CellPlex kit. 3 gene expression and associated multiplexing libraries were created with all 6 conditions being present in each library. Cells were sequenced through UCSF’s Center for Advanced Technology using NovaSeqX 10B.

#### Experiment 2

After tissue digestion of back skin (as above) and single cell solution formation, CD45+ and CD45-cells were sorted using in a 1:1 ratio from mock or *C. auris* colonized B6 mice, mock *C. auris* colonized IFNγ^-/-^ mice, *C. auris* colonized IFNγOE and *C. auris* colonized IL-17-/- mice. Prior to sorting 3-4 mice per condition were pooled. After sorting, 1 replicate per group was multiplexed in equal ratios using 10x 3’ CellPlex kit. 3 gene expression and associated multiplexing libraries were created with all 5 conditions being present in each library. Cells were sequenced through UCSF’s Center for Advanced Technology using NovaSeqX 10B.

### scRNAseq Data processing and analysis

Sequencing data were aligned to mouse genome mm10 with CellRanger version 8.1.0 (10x Genomics). Downstream data analysis, including clustering, visualizations and exploratory analyses was performed using the Seurat R package 5.0.2. Cells with <200 features, 6,500 reads (6,000 for experiment 2) or >6% mitochondrial genes were filtered out during preprocessing. The Seurat SCTransform function was used to normalize data using percent.mt as the variable to regress.

#### Experiment 1

Principle components were calculated using Seurat’s *RunPCA* function, followed by graph-based clustering using Seurat’s *FindNeighbors* (dims =1:10) and *FindClusters* functions and 2D visualization using Seurat’s *RunUMAP* function (dims = 1:10). Clusters were grouped based on cell-type defining markers (Fig1s2).

#### Lymphocytes

To define effector αβ T-cells, regulatory T-cells, effector γδ T-cells, dendritic epithelial T-cells (DETC), and innate lymphoid cells (ILCs), lymphocytes were reclustered twice. Using *RunPCA* function, *FindNeighbors* (dims=1:25), *FindClusters* functions and 2D visualization using Seurat’s *RunUMAP* function (dims=1:25) revealing Tregs, DETCs and Effector T-cells using cell type defining markers (Fig1s2C). Effector T-cells were reclustered (*RunPCA*, *FindNeighbors* (dims=1:20), *FindClusters*, *RunUMAP* function (dims=1:20)) revealing effector αβ T-cells, γδ T-cells and ILCs.

#### Myeloid cells

To define myeloid subsets we used *RunPCA* function, *FindNeighbors* (dims=1:25), *FindClusters* functions and 2D visualization using Seurat’s *RunUMAP* function (dims=1:25). Clusters were annotated using subset defining gene signatures (Fig2s3).

#### Keratinocytes

To define keratinocyte subsets we used *RunPCA* function, *FindNeighbors* (dims=1:25), *FindClusters* functions and 2D visualization using Seurat’s *RunUMAP* function (dims=1:25). Clusters were annotated using subset defining gene signatures (Fig3s1).

#### Experiment 2

After quality control as above, *RunPCA* function, followed by graph-based clustering using Seurat’s *FindNeighbors* (dims =1:10) and *FindClusters* functions and 2D visualization using Seurat’s *RunUMAP* function (dims = 1:10). Clusters were grouped based on cell-type defining markers (Fig4s4).

#### Keratinocytes

To define keratinocyte subsets we used *RunPCA* function, *FindNeighbors* (dims=1:15), *FindClusters* functions and 2D visualization using Seurat’s *RunUMAP* function (dims=1:15). Clusters were annotated using subset defining gene signatures (Fig3s1).

### scRNAseq Gene set enrichment analysis (GSEA) and over representation analysis (ORA)

In experiment 1, we used the ClusterProfiler package and used the gseGO and EnrichGO functions to perform GSEA and ORA respectively. Full gene ontology (GO) lists of upregulated genes using both approaches in *C. auris* colonized (compared *to* Mock and *C. albicans* colonized) and *C. albicans* colonized (compared to Mock and *C. auris* colonized) colonized mice are available in the supplementary materials. For graphical representation in Fig3s1 and Fig4s1, the Simplify function with Wang semantic similarity correction was used to remove redundant modules. In experiment 2, full GO lists of upregulated genes using ORA (EnrichGO) are available for C. auris colonized IFNγ^OE^ (compared to *C. auris* colonized B6 and IFNγ^-/-^ mice) and *C. auris* colonized IFNγ^-/-^ (compared to *C. auris* colonized B6 and IFNγ^OE^ mice).

### Gene module selection and statistical tests for DotPlots

For experiment 1, gene lists (TableS1) were curated following a literature search or selected from the ORA upregulated gene list (with adjusted p-value <0.05) in the associated GO term. For experiment 2, upregulated GO terms (adjusted p-value <0.05) were cross-referenced to GO terms upregulated in experiment 1 (*C. auris* colonized IFNγ^-/-^ upregulated GO terms were found in the *C. albicans* colonized B6 upregulated GO list and *C. auris* colonized IFNγ^OE^ upregulated GO terms were found in the *C. auris* colonized B6 GO list). For curated gene lists (i.e. epithelial migration^42^, necroptosis^43^, Cornified envelope, Tight junction, and Keratinocyte AMPs), statistical analysis was performed in R using either one way ANOVA (aov() function) with Tukey’s multiple comparison (TukeyHSD() function) test, student T-test (t.test() function), or Kruskal-Wallis test (kruskal.test() function) with Pairwise Wilcoxon Rank Sum Tests (pairwise.wilcox.test() function) for multiple comparisons depending on whether data appeared normally distributed.

## Notes

### Competing Interest Statement

The authors have declared no competing interest.

